# *OsRAD23a* negatively regulates salt tolerance and phosphorus uptake in rice

**DOI:** 10.64898/2026.08.27.747644

**Authors:** Shohei Oguro, Bilal Ahmad, Jaspinder Singh Dharni, Anil Kumar Nalini Chandran, Chi Zhang, Harkamal Walia

## Abstract

Salinity stress affects rice productivity due to reduced growth and sodium ion toxicity. Previously, we identified a splice variant of *RADIATION SENSITIVE23a* (*RAD23a*) as the potential basis for variation in salt-tolerance in rice germplasm. RAD23 is a known moonlighting protein associated with protein degradation. To validate the role of *RAD23a* in salt stress response, we characterized gene edited mutant lines that targeted the UBL and UBA2 domains of this protein. Mutation in either domain promoted shoot growth under saline and control conditions. The mutants also differed from wildtype plants in Na and K accumulation in roots and shoots under salt stress. Transcriptome analysis of mutants *versus* wildtype showed differential transcript abundance of multiple inorganic phosphate (Pi) starvation related genes, including *OsSPX2* and *OsPHO2*. As a result, mutants accumulate higher Pi compared to wildtype plants. The two allelic groups for *RAD23a* locus also differ in root and shoot phosphorus (P) content. Further, we show that RAD23a interacts with OsSPX2, a negative post-translational regulator of OsPHR2, the master regulator of Pi starvation response. Mutants have higher shoot growth and Pi levels under low Pi conditions, linking enhanced growth of mutants to increased Pi uptake. The UBA2 domain specific mutants have higher single grain weight and per plant grain weight than wildtype. In summary, we show that the *RAD23a* regulates differential growth, salt response and Pi uptake in rice in a domain-specific manner supporting the moonlighting roles of *RAD23a* in salt tolerance and phosphorus-dependent shoot growth.

## Introduction

Soil salinity is a major environmental stress that decreases crop productivity worldwide. Rice (*Oryza sativa*) is one of the most salt-sensitive, staple crops and exhibits a wide range of sensitivity to salt at different development stages (Zeng *et al*., 2002). It is moderately salt- tolerant during the germination, active tillering and maturation stages. It is highly susceptible during the early seedling and early reproductive stages (Munns and Tester, 2008). Plants under saline environment experience two major stresses, osmotic stress and ionic stress, which can ultimately lead to oxidative stress (Fu and Yang, 2023). Osmotic stress is caused by hyperosmotic soil conditions disrupting cell turgor and subsequently decreasing shoot growth rate during early phases of salt stress (Ponce *et al*., 2021). Ionic stress results from excessive accumulation of Na^+^ ions in plant cell cytoplasm, altering cellular ion balance, especially Na^+^ and K^+^ concentrations inside the cell and disrupting many biological processes (Munns and Tester, 2008). In natural soil conditions, ionic stress typically builds up over a longer duration and results in leaf senescence due to toxicity. Besides high level of sodium, salinity can also disrupt nutrient cycling in alkaline soils, especially phosphorus cycling, which lowers bioavailability of phosphorus (Bidalia *et al*., 2019; Ding *et al*., 2020).

Plants have evolved various mechanisms to cope with salt stress, among which high-affinity K^+^ transporters (HKTs) family has been extensively studied due to its potential role in Na^+^ homeostasis during salinity (Mian *et al*., 2011). HKTs are classified into two subgroups based on their phylogeny. Several group I HKT members (*OsHKT1;1*, *OsHKT1;4*, and *OsHKT1;5*) have functions of excluding excessive Na^+^ from xylem, thereby restricting accumulation of salt in leaf tissues (Hauser and Horie, 2010; Khan *et al*., 2020). Group II HKT including *OsHKT2;1* is associated with K^+^ transport as a Na^+^/K^+^ transporter (Hartley *et al*., 2020). Alternate splicing (AS) is another mechanism that is used by organisms to expand the range of proteome and regulate transcript abundance in response to abiotic stress including salinity stress (Sun *et al*., 2024; Zhou *et al*., 2021). Several salt-tolerance associated genes including *OsHKT1;4*, a sodium transporter linked to natural variation in shoot Na^+^ accumulation, is known to be alternatively spliced (Cotsaftis *et al*., 2012).

Excess Na^+^ accumulation under salt stress can disrupt the uptake, transport and accumulation of essential macronutrients, including nitrogen (N), phosphorus (P) and potassium (K^+^) (Iqbal *et al*., 2018; Razzaq *et al*., 2020). Phosphorus exists in organic and inorganic forms in soil. Plant can only assimilate inorganic phosphate (Pi), mainly as HPO_4_^2-^ and H_2_ PO_4_^-^, (Gho *et al*., 2020). Pi homeostasis is closely involved in salt stress mechanisms in plants (Ahmed *et al*., 2018; Hasanuzzaman and Fujita, 2022; Su *et al*., 2022). For instance, higher Pi accumulation in *Siz1* and *Pho2* mutants in Arabidopsis increased salt tolerance (Miura *et al*., 2011). While the interaction between Na^+^ and Pi remains poorly understood, numerous studies have demonstrated key components of the sensing and signaling systems involved in Pi starvation responses in rice (Gho *et al*., 2020). A key regulator in the Pi starvation response is a MYB transcription factor (TF), *Phosphate Starvation Response Regulator 2* (*OsPHR2*), a rice homolog of Arabidopsis *AtPHR1* (Wu *et al*., 2013). *OsPHR2* functions as central transcriptional activators for phosphate starvation-induced (PSI) genes responsible for Pi absorption and remobilization under Pi deficient conditions (Wu *et al*., 2013). These PSI genes harbor a cis-regulatory elements, P1BS, in the promoters that can directly bind to OsPHR2 and regulate the expression of miR399 and phosphate transporter family 1 (*OsPHT1*) genes (Lv *et al*., 2014; Zhou *et al*., 2008). Overexpression of *OsPHR2* resulted in increased Pi accumulation in shoots under the Pi-sufficient conditions (Liu *et al*., 2010; Zhou *et al*., 2008). It has been reported that SPX domain proteins negatively regulate OsPHR2 activity through physical interaction, which prevents binding of OsPHR2 to P1BS elements in PSI genes (Zhong *et al*., 2018). There are four SPX proteins in Arabidopsis and six SPX proteins in rice (OsSPX1-OsSPX6) (Duan *et al*., 2008). Furthermore, Gho et al. (2020) reported that overexpression of *OsPHR2* led to the upregulation of S-like RNase (RNS) family genes, belonging to class I and class II of RNase T2, which promoted RNA degradation and increased internal Pi levels under Pi starvation.

Genome-wide association analysis has been used to identify genetic loci regulating salt tolerance in rice (Campbell *et al*., 2017; Cui *et al*., 2024; Lv *et al*., 2022). With the advent of high-throughput plant phenotyping, integration of image-based phenotyping data with association studies has been investigated in response to salt stress in rice, aiming to capture the dynamics of stress responses in a nondestructive manner (Al-Tamimi *et al*., 2016; Campbell *et al*., 2015). Alternative splicing plays a critical role in gene regulation in eukaryotes as the variation in alternative splicing sites enables the production of various mRNAs variants from a single gene (Rosenkranz *et al*., 2022). A recent GWAS study integrated genome-scale mRNA splicing events as markers for association with shoot Na^+^ accumulation and shoot growth over time under salt stress to assess the impact of ionic stress and osmotic component of salt stress (Yu *et al*., 2021). One of the notable findings from this analysis was the discovery of a causal allelic variant (G/A) at the 6^th^ intron in *RADIATION SENSITIVE23a* (*RAD23a*) leading to presence or absence variation for an intron. The analysis showed that the minor (A) allele leads to intron retention and a premature stop codon (TAG), resulting in a truncated protein without C-terminus STI1 and UBA2 domains (Yu *et al*., 2021). In accessions with the major (G) allele, the protein is 392 amino acid (aa) long compared to 209 aa in the A allele accessions. The A allelic group of genotypes exhibited significantly higher Na^+^ concentration in the shoot than G allelic genotypes. Furthermore, an image-based shoot growth response for the diverse rice genotypes was used to obtain the projected shoot area (PSA) for the same set of rice accessions. This dataset was used to evaluate the impact of osmotic stress on shoot growth between *RAD23a* G/A alleles. The results showed a significant reduction in growth response under salt stress for the A allelic group relative to the G allelic group, suggesting that the splice variant in *RAD23a* could be potentially regulating variation in salt response in rice (Yu *et al*., 2021). However, the role of *RAD23a* as a candidate for regulating variation in salt tolerance in rice has not been shown.

RAD23 is a classical moonlighting protein that independently carries out multiple functions such as nucleotide excision repair (NER) and targeting protein degradation through the ubiquitin-proteasome pathway (Grønbæk-Thygesen *et al*., 2023). Rice has four RAD23 family members: *RAD23a* (*Os09g24200*), *RAD23b* (*Os02g08300*), *RAD23c* (*Os08g33340*), and *RAD23d* (*Os06g15360*) (Wang *et al*., 2020a). RAD23 proteins contain four distinct domains (Wang *et al*., 2017). One ubiquitin-like domain (UBL) located at the N-terminus interacts with 26S proteasomes. Two ubiquitin-associated domains (UBA1 and UBA2) are in the central and the C-terminus, respectively, and are known to interact with the ubiquitin or polyubiquitinated proteins. UBA2 domain is also known to be required for RAD23 stabilization to protect itself from proteasomal degradation in yeast (Heessen *et al*., 2005). Stress-inducible-1 domain (STI1) is localized between the two UBA domains, which contributes to DNA damage recognition (Grønbæk-Thygesen *et al*., 2023; Wang *et al*., 2017). UBL-UBA protein OsDSK2a mediates seedling growth and salt response in rice by binding with polyubiquitin chains and interacting with the gibberellin (GA)-deactivating enzyme ELONGATED UPPERMOST INTERNODE (EUI) resulting in its degradation through the ubiquitin-proteasome pathway (Wang *et al*., 2020a).

In this study, we tested the hypothesis that *RAD23a* regulates rice shoot growth in saline conditions, to determine the link between the splice variant predicted from the genome wide association analysis and salt tolerance. Since the A allele genotypes have a truncated protein, resulting in loss of some domains, we tested if the loss of specific domains of RAD23a between A and G allele genotypes may underlie the allelic difference in salt response. For this, we generated multiple gene edited lines and found a differential shoot growth response relative to wildtype plants. A transcriptomic analysis to gain molecular insights into the differential salt stress response showed a striking difference in transcript abundance of genes regulating inorganic phosphate (Pi) transport and signaling. We show that this transcript level differences in Pi-related genes resulted in difference in Pi uptake between the mutants and wildtype plants. Mechanistically, RAD23a interacts with OsSPX2, a negative post-translational regulator of PHR2, through co-immunoprecipitation and bimolecular fluorescence complementation assays. We also present evidence of domain specificity (UBL versus UBA) functions of the mutants for several of the phenotypic and transcriptomic differences that were observed between the wildtype and mutants, thus suggesting the moonlighting roles of *RAD23a* to extend to salt tolerance, and phosphorus uptake.

## Materials and Methods

### Plant materials and growth conditions

Rice seeds were surface-sterilized with 70% bleach (v/v) for 20 mins, soaked in sterile water overnight, and germinated on half-strength Murashige-Skoog (MS) agar for 2d in the dark, followed by 3d in the light. Germinated seedlings were transplanted to pots filled with 3.2 kg 75/25 mixture of completely dried Turface® MVP® and Pro-Mix BX (w/w). Each pot was daily watered with 150 mL. To standardize soil water availability across the plants, at 9 days after transplanting (DAT), each pot was watered to a uniform weight in order to maintain approximately 3600 mL of water in the soil. Salt stress treatment was applied in two steps of 35 mM each with 150 mL salt solution (840 mM NaCl:30.8 mM CaCl_2_) at 10 DAT and 13 DAT to reach 70 mM at 13 DAT. Control plants were watered with 150 mL until 13 DAT. At 11 DAT, pots were loaded onto the automated conveyor belt and plants were imaged daily until 27 DAT.

From 14 DAT, water levels were monitored and adjusted daily by the Scanalyzer 3D system (LemnaTec, Germany), an automated weight-based irrigation system to maintain the water status. Over the period of the experiments, greenhouse temperatures were set at 28°C during the day and 23°C during the night with day/night length of 16h/8h. At the end of the experiment, roots and shoots samples were collected for dry weight (DW) measurement, and Na, K quantification. For the phosphate deficiency experiment, germinated seedlings were transplanted to Profile® Greens Grade™ and grown hydroponically using half-strength Yoshida solution (Yoshida *et al*., 1976).

For the Pi-sufficient and low Pi conditions, the concentration of NaH_2_PO_4_ was adjusted to 200 μM and 10 μM, respectively (Lv *et al*., 2014; Wang *et al*., 2014). The hydroponic solution was adjusted daily to pH 5.5, and the nutrient solution was replaced every other day. Frozen samples of roots and shoots were collected at 14 DAT and 20 DAT for qPCR and Pi quantification, respectively.

### Vector construction and generation of transgenics

To generate CRISPR-Cas9 mutants, single-guide RNAs (sgRNAs) were designed using CRISPR-P 1.0 (http://crispr.hzau.edu.cn/CRISPR/) (Lei *et al*., 2014) targeting two different domains (UBA and UBL2) of the gene (Figure 1A) and cloned following the protocol described in Lowder et al. (2015). Single-guide RNAs synthesized as oligonucleotides with Eps3I (aka BsmBI) overhangs were cloned in the Eps3I-digested pENTR vector pYPQ141C, respectively. The entry clones with corresponding gRNAs were recombined with pANIC6B as a destination vector and pYPQ167 as a Cas9 donor using the LR Clonase II enzyme mix (Thermo Fisher Scientific). The destination constructs were then transformed into Agrobacterium tumefaciens strain EHA105, which were further used for rice transformation using calli of Kitaake, which has the major G allele for *RAD23a* splice variant (Yu *et al*., 2021). We identified and used only homozygous knockout, Cas9 free confirmed by β-glucuronidase screening assay, and T2 or later generations for all phenotypic and molecular analysis. The three independent events of *rad23a* mutants were referred to as mut1, mut2, and mut3 (Figure 1A).

**Figure 1.**
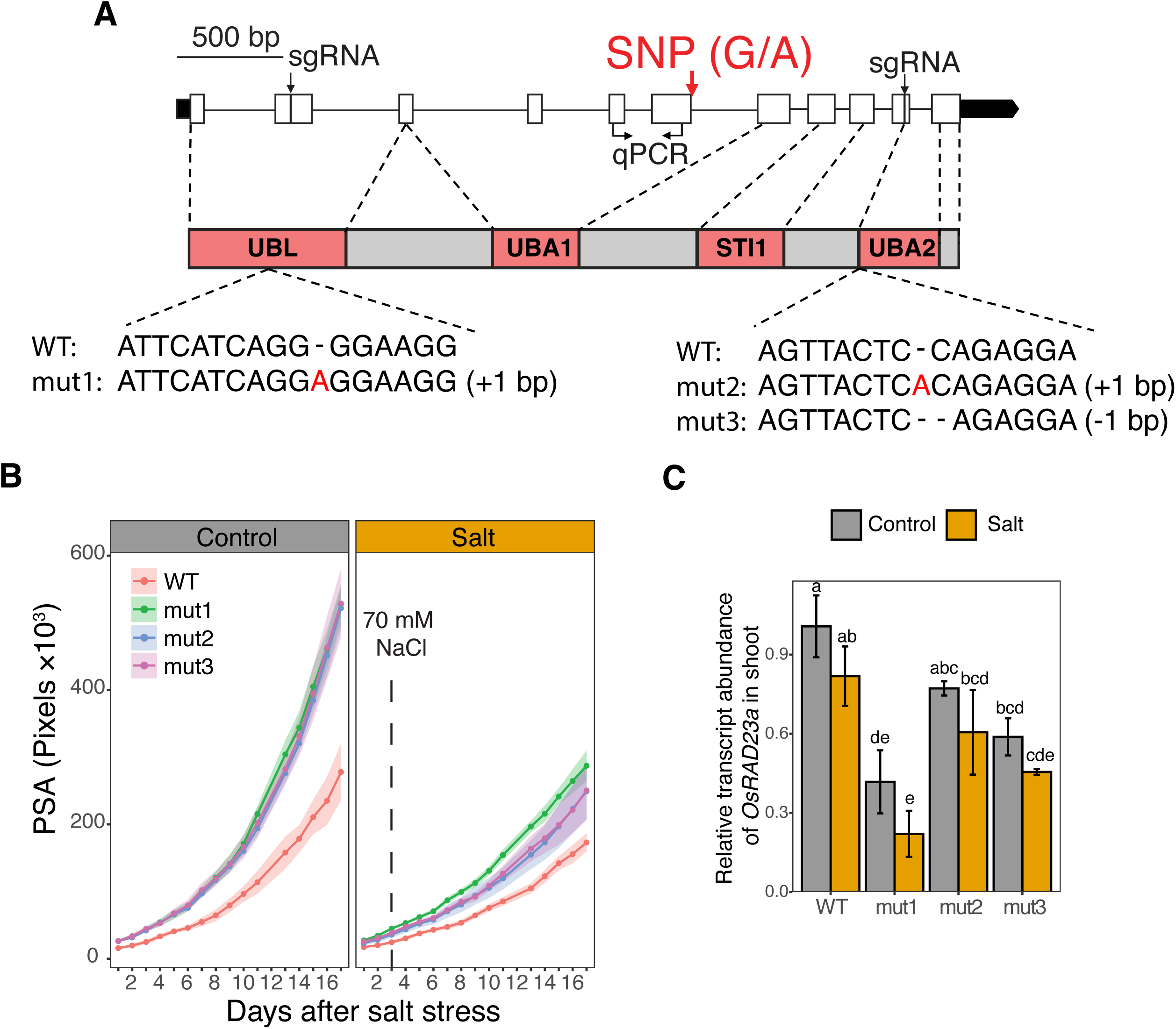
*RAD23a* gene model and image-based phenotyping growth response of *rad23a* mutants under control and salt stress for 17 days from the initiation of salt treatment. (A) The structure of the *RAD23a* gene and domain with the single nucleotide polymorphism (SNP) position at the first nucleotide in the 6^th^ intron and sgRNA. Untranslated region (black), exon (white), intron (solid line), and domain (red rectangle) were shown. (B) Projected shoot area (PSA) of wildtype (WT) and mutants (n=4). The solid line indicates the mean while the shaded area indicates the standard errors. A red vertical dashed line indicates the time point when 70 mM NaCl were applied. (C) Relative transcript abundance of *RAD23a* at 17 DAT, 7 days after the initial salt stress, in main shoot. Data represent means from two biological replicates, each with two technical replicates. Error bars represent ± SE. Statistical significance was determined using two-way ANOVA followed by Duncan’s multiple range test. Different letters denote significant differences at *P* < 0.05.

### RGB image capture and processing

Plants were imaged for 18 days after the initial salt application (11 DAT) until the end of experiment (27 DAT). For each plant, 10 side-view images, in which pot was rotated every 36° with exception of 90° instead of 180° (0°, 36°, 72°, 90°, 108°, 144°, 216°, 252°, 288°, and 324°), were captured every day. Due to the technical error in the imaging system on the 12 days after the initial salt stress, which is 22 DAT, the corresponding data is missing from the downstream analysis. Pixel counts of shoot regions were extracted from RGB side-view images using an open-source software PhenoImage (Zhu *et al*., 2021). All pixels from 10 RGB images were summed and defined as projected shoot area (PSA).

### Fluorescence image capture and processing

Fluorescence images were captured from 4 different side angles (0°, 72°, 144°, 216°) for each plant from 11 DAT to 27 DAT except 22 DAT. Plants were imaged under a continuous blue light (400-500 nm) and a fluorescent camera captured steady-state chlorophyll fluorescence from 500 to 700 nm. Fluorescence images were processed to obtain pixel counts and intensity using PhenoImage, where pixel counts were further categorized into 32 color classes (CC) based on image color ranges (Campbell *et al*., 2015; Zhu *et al*., 2021). Pixel counts from 4 side-view images were summed for each CC, and hierarchical cluster analysis (HCA) was performed to analyze the relationship between CC and time points across wildtype (WT) and mutants. CC with consistently low values were removed before performing the HCA.

### Quantification of Na, K concentration

Root and shoot samples were thoroughly washed with deionized water 3-4 times and dried at 75°C for at least 3d in an oven before obtaining dry weight. Oven-dried root and shoot samples were grinded into fine powder. Nitric acid (70%) was added into the samples (30 mg of root and 100 mg of shoot) to achieve a digest concentration of 100 mg/mL, and they were incubated overnight at 65°C for the digestion. The digests were cooled down to room temperature, centrifuged, and diluted 20-fold to a final concentration of 5 mg/mL dissolved solids. The Na, and K concentrations were quantified by inductively coupled plasma-mass spectrometry (ICP-MS, Agilent 7500cx), with a 2% nitric acid spiked with 50 ug/L Gallium as internal standard as described previously (Nývltová *et al*., 2022).

### Photosynthetic measurements

At 22 DAT, 12 days after the initial salt stress, the youngest fully expanded leaf of the primary tiller was measured (n=3) between 10:30 to 14:00 h to assess the physiological responses to salt stress in WT and mutants using the LI-6800 gas exchange system (LI-COR, USA). The chamber condition was maintained as follows: light intensity at 800 μmol m^-2^ s^-1^, reference CO_2_ concentration at 400 µmol mol^-1^, flow rate at 500 µmol s^-1^, relative humidity at 50%, and chamber pressure at 0.1 kPa. The evaluated photosynthetic parameters included CO_2_ assimilation rate and transpiration rate.

### Genomic DNA and RNA extraction and RT-qPCR

For *rad23a* mutant screening, genomic DNA was extracted from leaf tissue using the sucrose method (Berendzen *et al*., 2005). PCR genotyping was performed using KAPA3G Plant PCR Kit (Kapa Biosystems) and Sanger sequencing. Total RNA was extracted from the roots and shoots of control and salt stressed plants using Qiagen RNeasy Plant Mini Kit with DNase I digestion. Subsequently, 500 ng of total RNA were used for cDNA synthesis using iScript cDNA Synthesis Kit (Bio-Rad). Quantitative reverse transcription polymerase chain reaction (RT-qPCR) was performed using SYBR Green Master Mix (Roche) with the gene-specific primers. *OsOBP* (*Os03g16690*) and *OsUBQ5* (*Os01g22490*) were used as the internal control for shoots and roots, respectively (Soni *et al*., 2021). The relative expression was calculated using the 2^−ΔΔCq^ method (Livak and Schmittgen, 2001). Primers used in this study are listed in Supporting Table S1.

### RNA-seq analysis

For transcriptome analysis, main shoot at 17 DAT, 7 days after the initial salt stress, were harvested and snap frozen in liquid nitrogen. Frozen tissue was stored in -80°C freezer until further use. Quality and integrity of extracted total RNA were verified via fragment analyzer. RNA was then processed to prepare RNA-seq libraries with Kapa Hyper Stranded mRNA library kit (Roche). The libraries were quantitated by qPCR and sequenced for 101 cycles from single end of the fragments on a NovaSeq X Plus with V1.0 sequencing kits. Fastq files were generated and demultiplexed with the bcl2fastq v2.20 Conversion Software (Illumina). The 100-nucleotide single-end reads were mapped against the rice reference genome (RGAP v7.0; https://rice.uga.edu) using HISAT2 (Kim *et al*., 2019). The mapped reads in the exon regions were quantified using featureCounts (Liao *et al*., 2014). Differential expression analysis was performed using the DESeq2 package (Love *et al*., 2014). Read counts were normalized and genes with consistently low expression (<5) in all the samples were removed for the downstream analysis. The differentially expressed genes (DEGs) were determined using the criteria of |log_2_(fold change)| ≥ 0.58 and adjusted p value < 0.05, where log_2_(fold change) ≥ 0.58 and log_2_(fold change) ≤ −0.58 were considered as upregulated DEGs, and downregulated DEGs, respectively.

### Quantification of Pi content

Root and shoot tissues at 17 DAT (same as RNA-seq time point) from salt experiment and at 20 DAT from low Pi experiment were used for the quantification of Pi content using the phosphomolybdate colorimetric assay (Ames, 1966; Jain *et al*., 2007). Briefly, about 100 mg tissue grounded in liquid nitrogen was used for inorganic phosphate extraction in 1% acetic acid. The supernatant was mixed with ammonium molybdate and ascorbic acid and then assayed for Pi concentration by colorimetric assay at 820 nm.

### Yeast two-hybrid (Y2H) assays

The synthesized coding sequences of RAD23a and selected candidates (Supporting Table S2&S3), flanked by attL1 and attL2 sites, were cloned into pDEST22 prey and pDEST32 bait vectors, respectively, using the LR Clonase II enzyme mix (Thermo Fisher Scientific). The Y2H assay was performed as described earlier (Apprill *et al*., 2026). In brief, the prey and bait vectors were transformed into Y187 and Y2HGOLD strains using the standard LiAc method. The transformants were selected using selective media deficient in tryptophan (-Trp) and leucine (- Leu), respectively. Diploids were achieved by mating both strains overnight on YPDA media followed by selection on -Leu-Trp (-L-W) media. Interaction analysis was performed by testing the activity of various reporter genes, including *HIS3*, *AUR1-C*, and *MEL1*.

### Co-immunoprecipitation (Co-IP) assays

The synthesized RAD23a and OsSPX2 flanked with attL1 and attL2 sites were cloned into the pEarleyGate 104 vector in fusion with N-terminal Enhanced yellow fluorescent protein (EYFP) using the LR Clonase II. The OsSPX2 and OsPHR2 were cloned in fusion with N-terminal mScarlet-I in the pEarleyGate 104 vector containing mScarlet-I instead of EYFP using the LR Clonase II. The resultant vectors were sequence-verified by whole-plasmid sequencing (Eurofins Genomics), followed by transformation to Agrobacterium, GV3101. The transformants were grown overnight to an OD600 of 1. Cells were pelleted and resuspended in infiltration buffer (10 mM MgCl_2_, 10 mM MES [pH 5.6], and 150 µM acetosyringone) to a final OD600 of 0.4.

Various combinations of bait and prey were mixed with p19, a suppressor of RNA silencing, and injected onto the abaxial side of six-week-old *N. benthamiana* leaves using a 1 ml needleless syringe. The plants were kept in dark, humid conditions for 12 hours post-infiltration and then moved to long-day conditions with a 16h/8h light-dark regime. The expression was confirmed by observing the leaf disk under a fluorescent microscope, followed by sample collection, flash freezing, and storage at -80C. The Co-IP procedure was performed as described previously (Apprill *et al*., 2026). In short, the frozen samples were ground in liquid nitrogen to a fine powder, then homogenized in lysis buffer and incubated on ice for 30 minutes. Cell debris was removed by centrifuging the samples at least three times at 15,000 rpm for 10 minutes at 4 °C. The cleared lysate was subjected to Co-IP using the GFP-Trap magnetic agarose beads (ChromoTek). The bound proteins were washed 5-6 times to remove any nonspecific binding.

The proteins were eluted by boiling the beads in Laemmli buffer at 80 °C for 20 minutes. The proteins were resolved using the 4-20% TGX (Tris-Glycine eXtended) Gels and transferred to PVDF membrane. Blots were probed with rabbit anti-GFP (1:5000, Invitrogen, A-11122) and anti-RFP 419 (1:10000, Rockland, 600-401-379) primary and HRP-linked goat anti-rabbit (1:5000; Enzo, ADI-SAB-300) secondary antibodies. The chemiluminescence was detected by incubating blots with SuperSignal West Femto Maximum Sensitivity Substrate (Thermo Fisher Scientific).

### Bimolecular fluorescence complementation (BiFC) assay

The cDNA sequences of proteins used in the BiFC experiment were amplified with the primers listed in the Supporting Table S1, then gel purified and cloned into pDONR221 P3-P2 or P1-P4 entry vectors using the BP Clonase II Enzyme mix (Thermo Fisher Scientific). The coding sequences were subsequently transferred to the pBiFCt-2in1-NN vector (Grefen and Blatt, 2012), fused with the N- and C-terminal halves of mVenus in various combinations using the LR Clonase II enzyme mix. The resulting vectors were sequence-verified and transformed into Agrobacterium, followed by tobacco infiltration as described above. Plants were grown under long-day conditions for two days post-infiltration before fluorescence examination. Leaf discs were then collected and analyzed using confocal microscope (Nikon A1-NiE) to detect the fluorescence. Images were captured at 514 nm excitation and 522-554 nm emission for mVenus, 561 nm excitation and 570-620 nm emission for RFP, and 640 nm excitation and 663-738 nm emission for chlorophyll fluorescence.

## Results

### *RAD23a* negatively regulates rice shoot growth

We sought to characterize the role of *RAD23a* in salt stress response in context of shoot growth (osmotic component of salt stress). To this end, we generated three *RAD23a* gene edited lines in the wildtype background cultivar Kitaake using CRISPR/Cas9 system. Kitaake carries the more salt-tolerant G-allele, that lacks a pre-mature stop codon (Supporting Figure S1). RAD23a protein consists of four distinct domains: ubiquitin-like (UBL), ubiquitin-associated 1 (UBA1), stress-inducible-1 (STI1), and UBA2 domains (Wang *et al*., 2020a; Wang *et al*., 2017). Since the *RAD23a* SNP localizes to the middle of the coding sequence for the UBA1 domain, we aimed to assess how disrupting flanking functional domains affects *RAD23a* function. For this, we designed two editing constructs: one targeting the upstream region corresponding to the UBL domain and the other targeting the downstream region corresponding to the UBA2 domain. The mutant 1 (mut1) carries a 1-bp insertion in the upstream of the gene, affecting the sequence corresponding to the UBL domain and potentially all downstream domains including UBA1, STI1 and UBA2 due to the predicted frameshift whereas mut2 and mut3 have 1-bp insertion and deletion in the UBA2 domain, respectively (Figure 1A).

Our image-based phenotyping of a diverse set of rice lines was able to distinguish the A/G allelic groups based on averaged shoot growth response to salt stress. Since salt stress differentially affects shoot growth on a temporal scale, we performed an image-based shoot phenotyping experiment to determine if the wildtype Kitaake and the three mutant lines of *RAD23a* exhibit a differential shoot growth trajectory under control and salt stress conditions. We maintained a similar experimental set-up to that used for identifying shoot growth differences between the A and G-allele of *RAD23a*. To monitor the temporal growth response, plants were exposed to salt stress of 70 mM NaCl with corresponding control plants and were imaged daily for 17 d after the initiation of salt stress. Shoot RGB imaging captures osmotic (growth) component of the salt stress (Campbell *et al*., 2015). All three mutants exhibited increased shoot growth (measured as projected shoot area; PSA) compared with WT under control and salt stress conditions (Figure 1B). Under control conditions, all three mutant lines exhibited the higher PSA compared to WT. Under salt stress, mut1 showed the highest PSA followed by mut2 and mut3 and with WT exhibiting the lowest PSA. Our imaging derived PSA and destructive shoot DW at 27 DAT, 17 days after the initial salt stress shows a strong positive correlation (*R* = 0.97, *P* < 2.2 × 10^-16^) supporting the accuracy of image analysis (Supporting Figure S2). At 27 DAT, all mutant lines also showed a significant increase in root biomass relative to WT under control whereas only a slight increase was observed under salt stress (Supporting Figure S3). Collectively, these data indicate that lines edited for *RAD23a* have higher shoot growth under control conditions relative to WT. Although the mutants exhibit sensitivity to salt stress, the shoot biomass is still higher than the WT. The salt-sensitivity of the individual mutants varies slightly, especially for mut1 (UBL domain), which is less salt-sensitive for shoot growth response than mut2 and mut3 (UBA2 domain). This indicates that *RAD23a* negatively regulates shoot growth.

We next examined the expression of *RAD23a* in the edited lines relative to the WT under control and salt stress conditions. For this, we measured the expression of WT and mutant lines at 17 DAT, 7 days after the salt stress. We found that the edited lines have slightly lower transcript abundance relative to WT plants (Figure 1C). There is variation in transcript abundance among the mutant lines with mut1 having the lowest expression. Transcript abundance of *RAD23a* is slightly reduced under salt stress in mutant and WT plants. This analysis suggests that the edit targeting the UBL domain (mut1) affected the *RAD23a* transcript abundance/stability more severely than the edits in the UBA2 domain in mut2 and mut3. Therefore, differences observed among the mutants could be due to disruption of the targeted domain and/or due to differences in transcript abundance.

### *RAD23a* regulates the ionic response under salt stress

We next examined the ion uptake and ion homeostasis in relation to higher shoot growth in *rad23a* mutants under both control and salt conditions. We measured the concentrations of Na, K in roots and shoots on the last day of experiment (27 DAT). Under salt conditions, Na concentration in roots was significantly reduced in mut2 and mut3 while shoot Na concentration decreased in mut1 and mut3 (Figure 2A). Mutants generally accumulated higher levels of K in roots but exhibited lower levels in shoots, regardless of treatment conditions (Supporting Figure S4A). Specifically, root K concentration was significantly higher in all mutants compared to WT under control conditions. Under salt stress, mut2 and mut3 maintained significantly elevated root K concentration. In shoots, mut3 accumulated less K than WT under control conditions, whereas mut1 and mut2 showed comparable K levels to WT. However, under salt stress, mut1 and mut2 have lower shoot K concentration. In some species, the Na/K ratio is considered as a more relevant trait associating with salt tolerance as Na interferes with K homeostasis (Shabala and Pottosin, 2014). Because of differences in Na and K concentration between WT and mutants, mut2 and mut3 maintained a significantly lower root Na/K ratio compared to WT under salt stress (Supporting Figure S4B). Similarly, shoot Na/K ratio was significantly decreased in mut1 and mut3 under salt conditions. This data indicates that root and shoot Na concentration for the mutants differed from wildtype plant in most cases but was not consistent across all three mutants.

**Figure 2.**
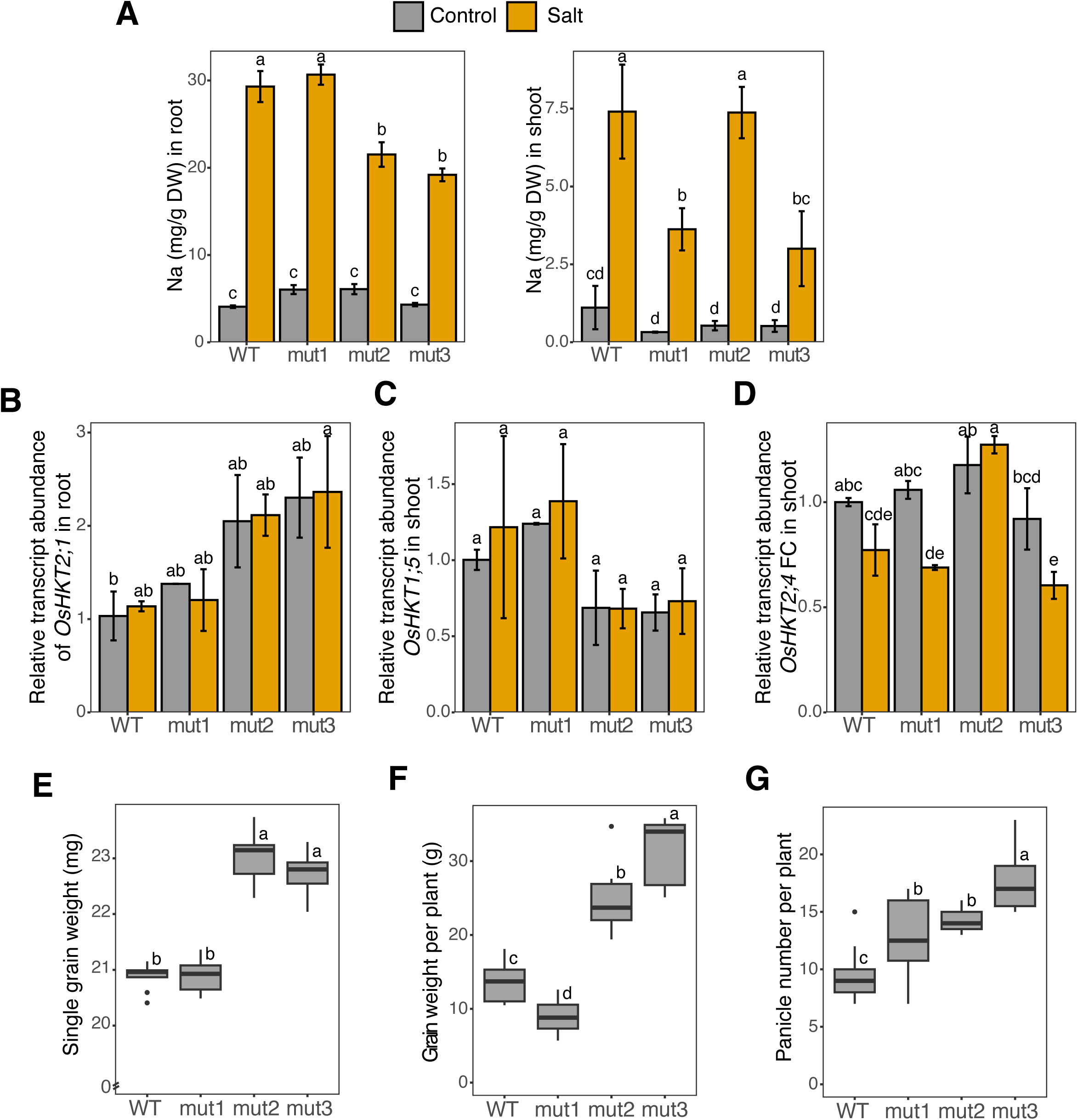
*rad23a* mutants accumulate lower sodium. (A) Sodium concentrations in roots and shoots under control and salt stress conditions at 27 DAT. (B-D) Transcript abundance of *OsHKT2;1* in roots, *OsHKT1;5* and *OsHKT2;4* in shoots under control and salt stress conditions at 17 DAT. Error bars represent ± SE. Statistical significance was determined using one-way or two-way ANOVA followed by Duncan’s multiple range test (n=3-4 for elemental analysis, n=2 with two technical replicates for expression analysis). Different letters denote significant differences at *P* < 0.05. (E) Single grain weight (mg) of mature grains grown in soil. Single grain weight was calculated from 50 grains per plant as a biological replicate. (F) Grain weight per plant (g). (G) Panicle number per plant. For yield-related traits (E-G), statistical significance was determined using one-way ANOVA followed by Duncan’s multiple range test (n=7-10).

Sodium ion homeostasis in plants is associated with a family of ion transporters called HKTs (Riedelsberger *et al*., 2021). We measured the expression level of seven *HKT* genes in shoots and roots at 17 DAT (Figure 2B-D; Supporting Figure S5). Notable among these, the expression of *OsHKT2;1* was higher in mut2 and mut3 in roots compared to WT under control and salt stress, but the differences were not statistically significant within each treatment (Figure 2B). *OsHKT1;5* mediates sodium exclusion from the xylem in roots to prevent its accumulation in leaf blades (Kobayashi *et al*., 2017). In shoots, *OsHKT1;5* expression did not differ significantly among genotypes, however, mut2 and mut3 expression level was slightly lower relative to WT (Figure 2C). Expression of *OsHKT2;4* was significantly higher in mut2 compared to WT under salt stress (Figure 2D). Collectively, these results suggest that *rad23a* mutants in Kitaake maintain a more favorable ionic status than WT under salt stress during vegetative development, but the expression level differences for the examined HKT family members did not establish a consistent link with sodium level differences.

As the mutants exhibited an increase in shoot growth, we then asked if the increased biomass at the vegetative stages translates into agronomic traits such as yield parameters. For that, we analyzed grain weight per plant and single grain weight from mature plants grown in field soil without any stress treatment. The UBA2 mutants (mut2 and mut3) significantly increased single grain weight compared to the WT seeds while UBL mutant (mut1) seeds were comparable as WT seeds (Figure 2E). The panicle number per plant was increased in all mutants, however, total seed weight per plant was higher only in UBA2 mutants but lower in UBL mutant relative to WT (Figure 2F&G). The decrease in panicle weight in mut1 may be associated with increased seed sterility. These results suggest that edits in UBA2 domain of *RAD23a* improves shoot growth and positively influences seed weight at maturity.

### *rad23a* mutants maintain higher photosynthetic rate under salt

Increased salinity negatively affects the physiological status of the plant especially photosynthesis. We evaluated the impact of edits made to *RAD23a*, by estimating the rate of leaf senescence, using a fluorescence camera and the same set-up used for shoot growth imaging under control and salt stress for 17 days. Fluorescence imaging captures the ionic component of salt stress due to degradation of photosynthetic pigments in response to increased Na concentration (Campbell *et al*., 2015). Fluorescence images were processed to obtain pixel counts and intensity, where pixel counts were further categorized into 32 color classes (CC) based on image color range (Campbell *et al*., 2015; Zhu *et al*., 2021). Hierarchical cluster (HCA) analysis showed that there are developmental, genotypic, and treatment-based differences in fluorescence pixel counts (Figure 3A). Specifically, for color class labeled as CC8, both WT and *rad23a* mutants were high during the early time points from 11 to 17 DAT, corresponding to 1 to 7 days after the initial salt stress. However, WT exhibited a gradual decline starting at 18 DAT with further reductions from 20 to 27 DAT under both control and salt conditions. In contrast, mut1, mut2 and mut3 exhibited consistently high pixel counts for CC8 over the 17d of imaging. For CC10, WT and mutant lines were similar from 21 to 27 DAT under control conditions, however, WT showed higher pixel count populating CC10 compared with mutants under salt conditions over the 17 days, as well as from 11 to 19 DAT under control. These results suggest that the mutation in *RAD23a* alters the dynamics of fluorescence signal (a proxy for chlorophyll pigmentation) under both treatments.

**Figure 3.**
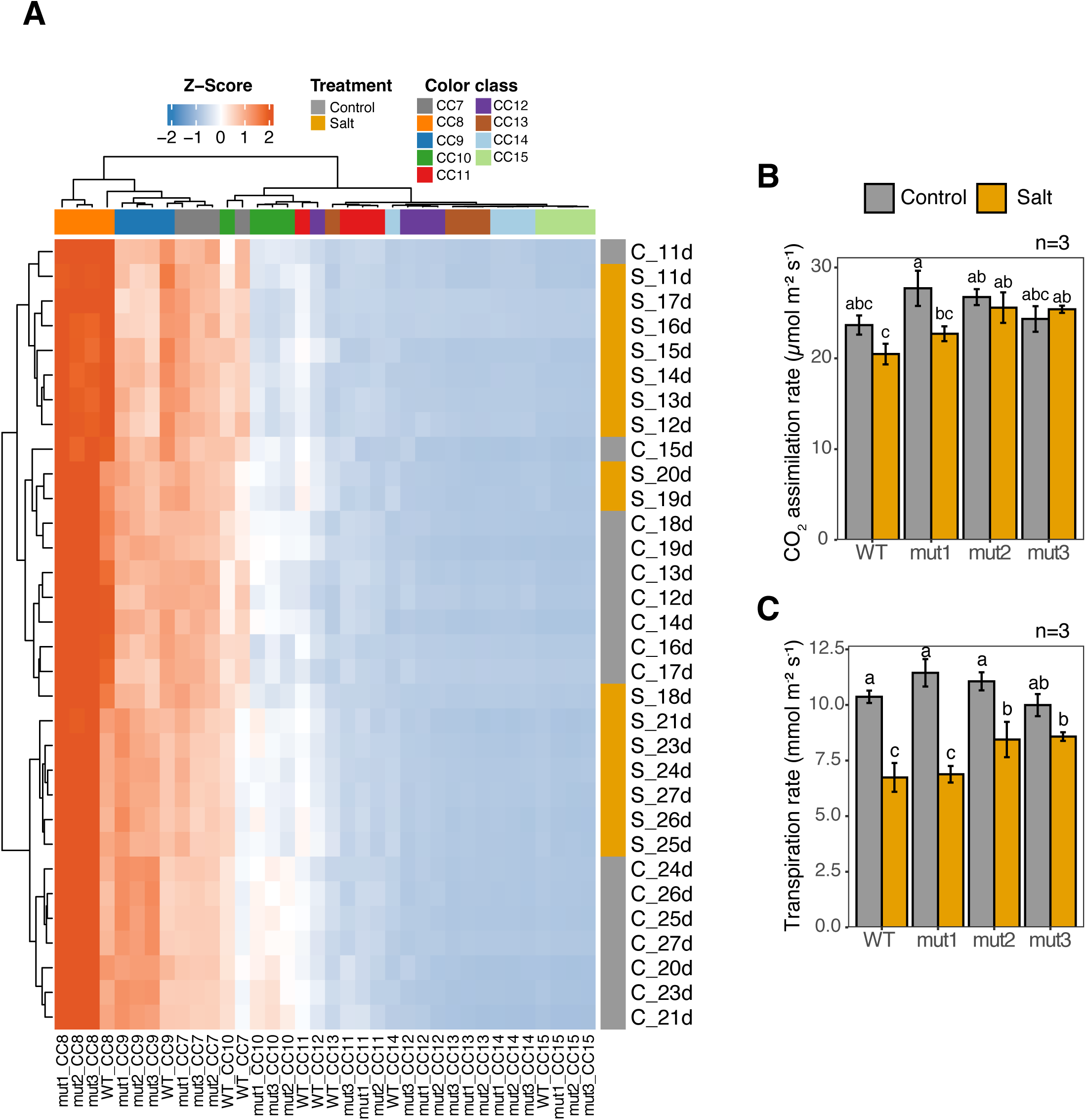
Clustering analysis of image-based fluorescent traits (A). Fluorescent pixel counts were categorized into different color classes (CC) based on image color ranges (Zhu *et al*., 2021; Campbell *et al*., 2015). On the X-axis, treatments are shown along with the data collection day in DAT. Photosynthetic parameters of leaves for WT and mutants under control and salt stress conditions at 22 DAT. (B) CO_2_ assimilation rate and (C) Transpiration rate. Error bars represent ± SE. Statistical significance was determined using two-way ANOVA followed by Duncan’s multiple range test (n=3). Different letters denote significant differences at *P* < 0.05.

Given the differences observed between the WT and mutants for pigmentation estimated through fluorescence intensity during the early stage (11 to 20 DAT) of imaging, we next sought to determine if these differences in pigmentation or chlorophyll manifest as changes in photosynthetic rate. For this, we measured photosynthetic parameters on leaves at 22 DAT under both control and salt stress conditions. There was no significant difference in CO_2_ assimilation rate between WT and *rad23a* mutants under control conditions (Figure 3B). Salt treatment for most genotypes reduced CO_2_ assimilation rate. However, mut2 and mut3 maintained a significantly higher level of CO_2_ assimilation rate than WT under salt stress. Transpiration rate was similar between WT and mutants under control conditions, but salt stress led to a significant decrease in transpiration rate in WT, mut1, and mut2, with no decline observed in mut3 (Figure 3C). Overall, the mut2 and mut3 maintained higher assimilation rate and transpiration rates than WT under salt conditions.

### *rad23a* mutants suppress induction of numerous salt responsive genes

To gain molecular insights into the differential growth and ionic responses of mutants relative to WT plants, we performed a transcriptome analysis for shoot tissue grown under control and salt stress conditions (imposed for 7 d) during early vegetative stage. We performed pair-wise comparisons for each mutant with WT plants to identify differentially expressed genes (DEGs). These comparisons yielded 206 and 97 genes that were upregulated and downregulated, respectively under control conditions. Under salt stress, 553 genes were upregulated, and 513 genes were downregulated. After removing the redundant DEGs from these two lists, we performed clustering analysis of 1,197 DEGs with the number of clusters (k) set to 15 (figure 4A). Two main transcriptomic patterns emerged from this analysis when represented as a heat map. Four of the fifteen clusters (Cluster: 4, 3, 1 and 2) exhibit an increase in transcript abundance of genes in WT in response to salt stress but lack a similar increase in mutants under salt stress. This suggests that salt stress imposed for this experiment upregulates a large number of genes (354 genes) in WT plants but edits/mutations in *RAD23a* suppress a major portion of the transcriptional response from manifesting or the increased salt tolerance of the mutants results in these plants not reaching the stress threshold at which these genes are upregulated significantly. This pattern is not evident for clusters populated by genes that are downregulated under salt stress in WT plants. The second transcriptomic pattern shows that the three mutants also have differentially regulated genes with transcript abundance that is specific to each of the mutants and different from WT. For instance, for mut1, Cluster 10 and 11 is predominantly populated by genes that have higher transcript abundance in control and salt stress conditions relative to all other genotypes. It is noteworthy that transcript abundance of *RAD23a* in mut1 is ∼2-fold lower relative to WT and the other two mutants in both control and salt stress treated plants (Figure 1C). Most genes in Cluster 12 have relatively higher expression in mut3 under control and salt stress conditions whereas genes in Cluster 13 exhibit higher transcript abundance in mut3 only under salt stress conditions.

**Figure 4.**
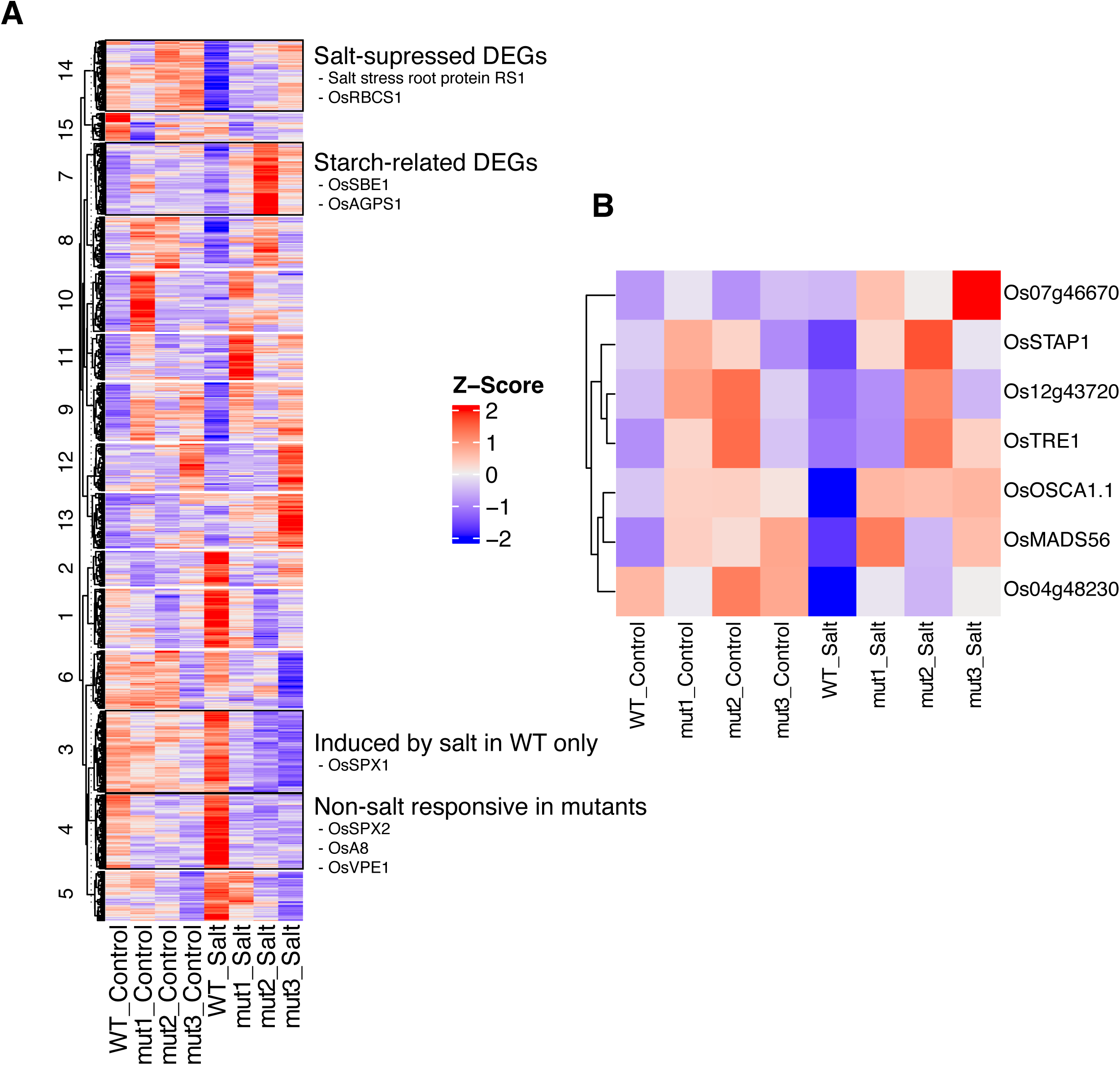
Transcriptome analysis and gene expression analysis under control and salt stress. (A) All the differentially expressed genes (DEGs) from pairwise comparison of each mutant and WT were shown. (B) Heat map of several salt- and/or dehydration-related DEGs.

Among stress-responsive pathways, multiple salt- and/or dehydration- related genes showed distinct regulation patterns between WT and mutants (Figure 4B; Supporting Table S4). *OsMADS56* was generally upregulated in mutants relative to WT under control and salt stress conditions. *OsMADS56* was identified as a causal candidate gene of salt tolerance associated with presence-absence variations and known to be involved in drought tolerance (Cui *et al*., 2024; Nurdiani *et al*., 2025). Expression of *OsTRE1* (*trehalase 1*) was higher in mut2 and mut3 relative to WT under salt stress. Overexpression of *OsTRE1* is known to enhance salt tolerance (Islam *et al*., 2019). Expression of *OsOSCA1.1* was decreased in WT under salt stress, while it was maintained or slightly increased in mutants. *OsOSCA1.1* is associated with stomatal closure and plant survival in response to hyperosmolarity and salt stress, and overexpression of *OsOSCA1.1* increased drought and salt stress tolerance (Han *et al*., 2022). Transcript abundance of *OsSTAP1* (an AP2/ERF-type transcription factor), positive regulator of decreasing Na^+^/K^+^ ratio (Wang *et al*., 2020b), was highly upregulated in mut2 relative to WT under salt stress.

We next mined the allelic variation in expression of seven selected salt- and/or dehydration-related DEGs using the RNA-seq dataset of rice diversity panel which was previously used for identification of *RAD23a* under salt stress (Yu *et al*., 2021). Among them, expression of *OsMADS56* and *OsOSCA1.1* exhibited the strong allelic variation in both control and salt stress conditions (Supporting Figure S6). The G allele genotypes showed several hundred-fold higher expression of *OsMADS56* relative to the A-allele genotypes. However, none of the seven examined genes showed the similar expression pattern as *RAD23a* where its expression in the A- allelic genotypes was significantly higher than the G-allelic genotypes under control and salt stress conditions (Supporting Figure S7).

### Differential regulation of inorganic phosphate related genes in *rad23a* mutants

One of the main pathway-specific features to emerge from the transcriptome analysis is the differential regulation of genes associated with the uptake and transport of inorganic phosphate (Pi) between WT and one or more of the mutant plants (Figure 5A). We selected this subset of twelve Pi-related genes and performed a cluster analysis. Among these genes, *SPX* [named after SYG1 (suppressor of yeast gpa1), Pho81 (CDK inhibitor in yeast PHO pathway), and XPR1 (xenotropic and polytropic retrovirus receptor)] (Wang *et al*., 2014) including *OsSPX1*, *OsSPX2*, and *OsSPX4* were generally downregulated in the mutants, with *OsSPX2* consistently showing lower expression across all three mutant lines relative to WT. In rice, OsSPX1 and OsSPX2 inhibit phosphate starvation responses in interaction with OsPHR2, a key regulator of phosphate starvation signaling (Wang *et al*., 2014). In contrast, *OsPHO2* (a ubiquitin conjugating E2 enzyme), that negatively regulates the inorganic phosphate starvation response, has higher transcript abundance in the mutants under both control and salt stress conditions. Similarly, S-like RNase (RNS) family genes *OsRNS4* and *OsRNS5* are also expressed more strongly in the mutants in both conditions. *OsRNS4* and *OsRNS5* are suggested to be involved in Pi accumulation, likely via negative regulation by *OsPHO2* and positive regulation by *OsPHR2*, based on expression analyses (Gho *et al*., 2020). Additionally, there was a significant *OsRNS4* expression difference between *RAD23a* A-allele and G-allele genotypes under salt stress conditions (Supporting Figure S8). Expression of *OsTRXh1* (an h-type thioredoxin) was upregulated in WT under salt stress relative to mut2 and mut3 but not under control condition. It has been reported that *OsTRXh1* transcript expression is induced by salt stress, and its overexpression causes salt sensitivity (Zhang *et al*., 2011). Further, OsTRXh1 interacts with OsPHO2, and suppression of *OsTRXh1* leads to a slight increase in Pi concentration (Ying *et al*., 2017). Notable among these genes is the expression of *OsPHT1;11*, which is specifically upregulated in mut1 in both control and salt stress treatment. *OsPHT1;11* is involved in symbiotic Pi absorption in rice (Yang et al., 2012). Expression of three additional Pi transporters, *OsPHT1;4*, *OsPHT2;1*, and *OsPHO1;3* is higher in WT under control and salt stress compared to most mutant samples. Collectively, the differential expression of Pi related genes between WT and mutants in both control and salt stress samples suggests that the *RAD23a* edits are likely altering the phosphorus status of the plants in a salinity-independent manner.

**Figure 5.**
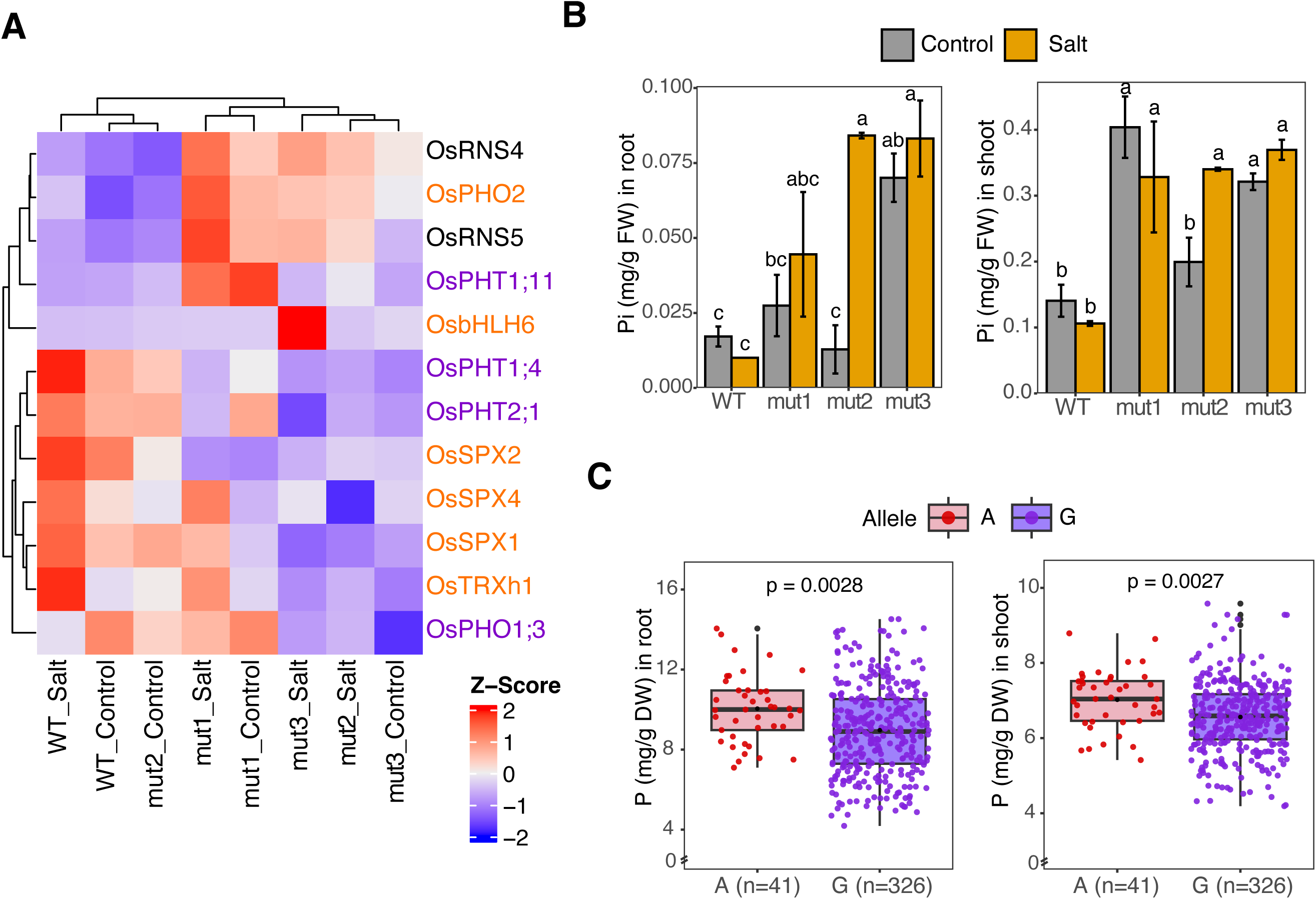
*rad23a* mutants accumulate higher amount of inorganic phosphate (Pi). (A) Heatmap of several Pi-related DEGs. Genes involved in Pi signaling and Pi transport are colored orange and purple, respectively. (B) Pi concentration in roots and shoots under control and salt stress conditions at 17 DAT. Statistical significance was determined using two-way ANOVA followed by Duncan’s multiple range test (n=3). Different letters denote significant differences at *P* < 0.05. (C) Allelic difference in P concentration in roots and shoots. A-allelic (n=41) and G-allelic (n=326) genotypes were compared using Students t-test.

We next tested if the transcript-level perturbation of phosphorus-related genes in *rad23a* mutants results in a change in phosphorus levels of the mutants. For this, we used the root and shoot tissues from the same timepoint and experiment from transcriptome analysis sampling to determine the inorganic phosphate (Pi) level. Inorganic phosphate levels were generally higher in mutants than the WT plants under control conditions and increased even more under salt stress (Figure 5B). It is noteworthy that salt stress increases the Pi concentration, and the genotypic differences are more evident in the salt stress samples. Further, our analysis indicates that there are Pi level differences among the mutants as well. For instance, shoots of mut1 under control conditions have higher Pi concentration than any other samples. Collectively, the differential expression of phosphorus-associated genes in shoot tissue and measured differences in Pi concentration between WT and mutant plants indicate that *RAD23a* is involved in determining phosphorus status in rice.

Since *RAD23a* as a locus exhibit differential salinity tolerance, we next asked if the natural variation at the SNP-9.14364417 (basis of A/G allele) within *RAD23a* also exhibits variation in phosphorus content in rice germplasm. To examine this, we used the publicly available ionome dataset for the rice diversity panel 1 (RDP1) by Cobb et al. (2021), where a subset of germplasm was used for identification of *RAD23a* alleles. We found that the minor A allele genotypes have significantly higher P concentration relative to the major G allele genotypes in the roots and shoots (Figure 5C). This indicates that natural variation at the *RAD23a* locus in rice could be potentially involved in regulating phosphorus uptake.

Since changes in phosphorus availability and uptake can significantly affect plant growth (Malhotra *et al*., 2018; Schachtman *et al*., 1998), we next asked if the difference in shoot growth observed between the WT and mutant plants persists when these genotypes are grown under Pi limiting conditions. For this, we imposed a low Pi treatment and normal Pi (Pi-sufficient) treatment in a hydroponics setup. Both under Pi-sufficient and low Pi conditions, all the edited lines showed significantly higher shoot DW and root DW relative to WT at 14 days after treatment (Figure 6A). We then determined Pi status in root and shoot tissues at 21 days after treatment (Figure 6B). Under Pi-sufficient conditions, mut1 significantly increased Pi level in roots compared to WT and other two mutants while under low Pi conditions, all the mutants exhibited higher Pi concentration in roots relative to WT. Notably, in the roots, Pi concentration in mut1 is nearly double the levels found in WT and other two mutants regardless of the treatment. In shoot tissue, all the mutants have significantly higher Pi concentration relative to WT. As in root tissue, mut1 maintains higher Pi concentration in shoot tissue relative to the other two mutants in both Pi-sufficient and low Pi conditions. There was no significant treatment effect on Pi level in shoot tissue within the genotype. Collectively, these data show that mutations in *RAD23a* enable the plants to maintain higher levels of Pi in shoot tissue and the specificity of the site of mutation can further augment this Pi levels in shoot as well as roots as evident from mut1.

**Figure 6.**
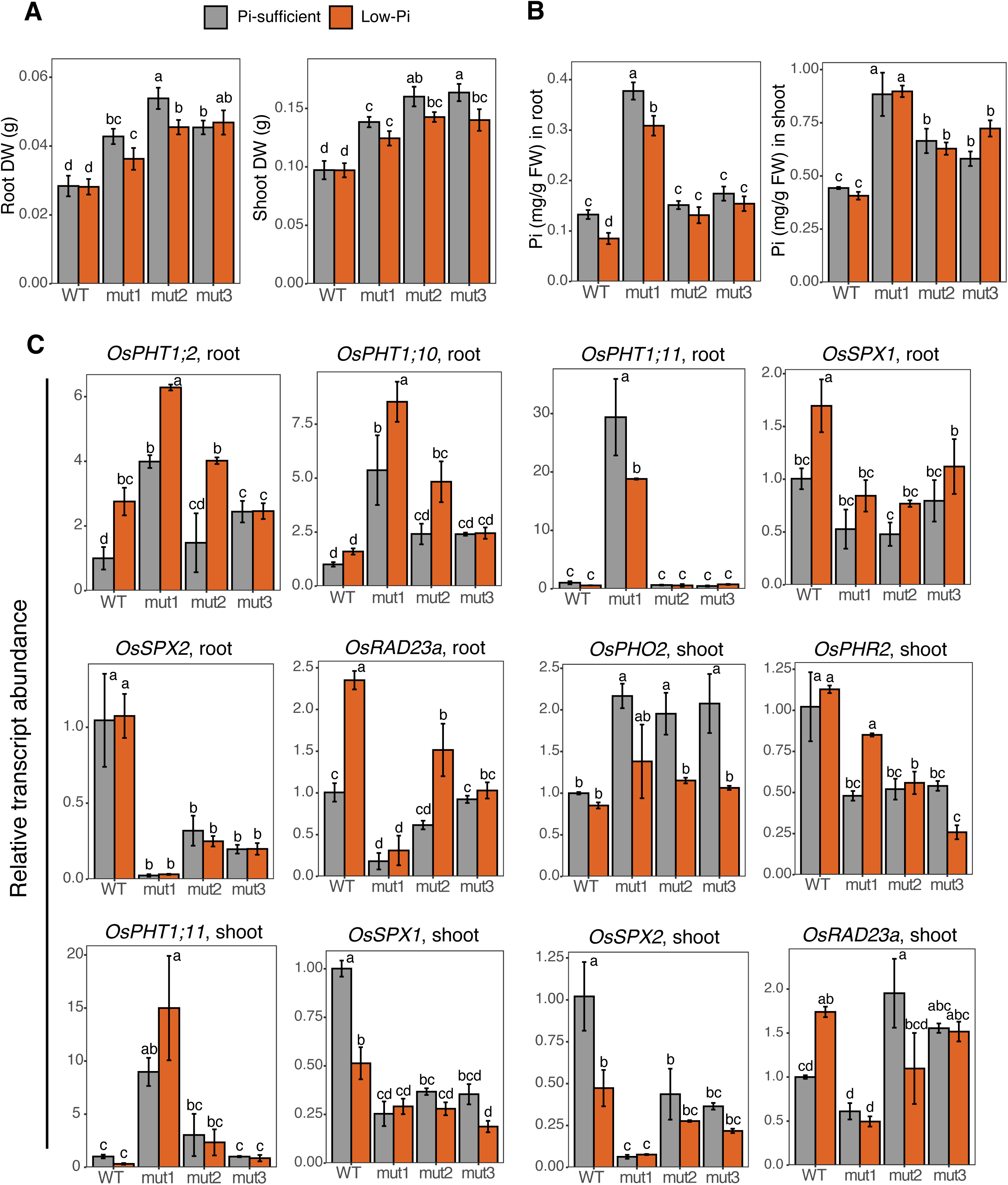
*rad23a* mutants exhibit higher root and shoot biomass with increased Pi concentration under Pi-sufficient and low Pi conditions. (A) Root and shoot dry weight (g) at 14 DAT, (B) Pi concentration in roots and shoots under Pi-sufficient and low Pi conditions at 20 DAT. (C) Relative transcript abundance of *RAD23a* and several Pi related genes at 14 DAT in roots and shoots. Statistical significance was determined using two-way ANOVA followed by Duncan’s multiple range test (n=5 for A and B, and n=2 for C). Different letters denote significant differences at *P* < 0.05.

We next performed expression (RT-qPCR) analysis to determine if low Pi treatment affects *RAD23a* transcript abundance. We found that *RAD23a* expression is upregulated in both root and shoot tissue of WT in response to low Pi treatment (Figure 6C). Among the mutants, only mut2 showed an upregulation in roots and downregulation in shoot tissue in response to low Pi treatment. Expression of *RAD23a* in mut1 in both tissues is lower than other genotypes consistent with the previous observation (Figure 6C; Figure 1C). We then assayed the expression of Pi-related genes including the DEGs identified from the transcriptome analysis for their response to the low Pi treatment (Figure 6C). Transcript abundance of *OsSPX1* and *OsSPX2* is significantly higher in WT relative to the mutants under low Pi treatment in roots. Expression of *OsSPX1* increased in WT roots in response to low Pi while *OsSPX2* did not respond to low Pi treatment in roots of any genotype. The shoot transcript level response of *OsSPX1* in WT was opposite of that in roots, as it decreased in response to low Pi treatment. The transcript abundance of *OsSPX2* in shoots also decreased in WT but did not change significantly in the mutants. To summarize, *OsSPX1* and *OsSPX2* exhibit tissue specificity in response to low Pi treatment in WT, but their transcript abundance does not change significantly in response to low Pi in roots or shoots in the mutants. The expression of *OsPHR2* was decreased in the mutants relative to the WT in shoot tissue under Pi-sufficient and low Pi conditions. The expression of phosphate transporters *OsPHT1;2* and *OsPHT1;10* was increased in root tissue in response to low Pi treatment, and both transcripts were generally upregulated in the mutants relative to WT under Pi-sufficient and low Pi treatment (Figure 6C). Another notable genotypic difference was significantly higher *OsPHO2* expression in mutants relative to WT in shoot under Pi-sufficient conditions. Under low Pi, the mutants exhibited a much lower *OsPHO2* transcript level relative to Pi-sufficient condition but comparable to WT plants. Further, consistent with the transcriptomic data, we observed a striking induction of *OsPHT1;11* in mut1 in root and shoot tissue under both conditions relative to WT and the other two mutants. Collectively, these results suggest that the mutation in *RAD23a* increases the expression of phosphate transporters by reducing the *OsSPX2* transcript abundance, thereby promoting Pi uptake.

### RAD23a interacts with OsSPX2 but does not trigger its turnover

To dissect the molecular mechanisms of RAD23a-mediated Pi regulation, we investigated its interactions with several candidate proteins selected from Pi related DEGs identified in the transcriptomic analysis, including OsSPX1, OsSPX2, OsPHO2, OsRNS4, and OsRNS5. We also included OsPHR2 in the assay as it is a key regulator of the Pi starvation response, although it was not differentially expressed. First, we performed yeast two-hybrid (Y2H) assays for direct interaction between RAD23a and these proteins. We did not observe a direct interaction between RAD23a and the selected seven candidates (Supporting Table S2). Then, we tested indirect interaction between RAD23a and six selected candidates, namely OsSPX1, OsSPX2, OsPHR2, OsPHO2, OsRNS4, and OsRNS5. For the indirect interaction assay, we fused the proteins with EYFP or mScarlet-I and transiently expressed them in *N. benthamiana,* followed by Co-IP. Notably, we found an interaction between RAD23a and OsSPX2 (Figure 7A). However, we failed to detect an interaction between RAD23a and OsSPX1. Similarly, a very weak to no interaction was observed with OsPHR2. Since RAD23a is a UBA-UBL domain containing protein and is known to play a role in 26S proteasomal degradation (Tsuchiya *et al*., 2017; Zhang *et al*., 2009), we hypothesized that it might affect OsSPX2 accumulation. To this end, we expressed OsSPX2 with EYFP alone and EYFP fused to RAD23a (EYFP:RAD23a). Additionally, we included a replicate treated for 8 h with MG132 before sample collection to observe any changes in OsSPX2 accumulation. However, we did not find any changes in the OsSPX2 levels in the presence and absence of MG132 and RAD23a in tobacco (Figure 7A). This implies that RAD23a interact with OsSPX2 but does not target OsSPX2 for its turnover through the 26S proteasome pathway.

**Figure 7.**
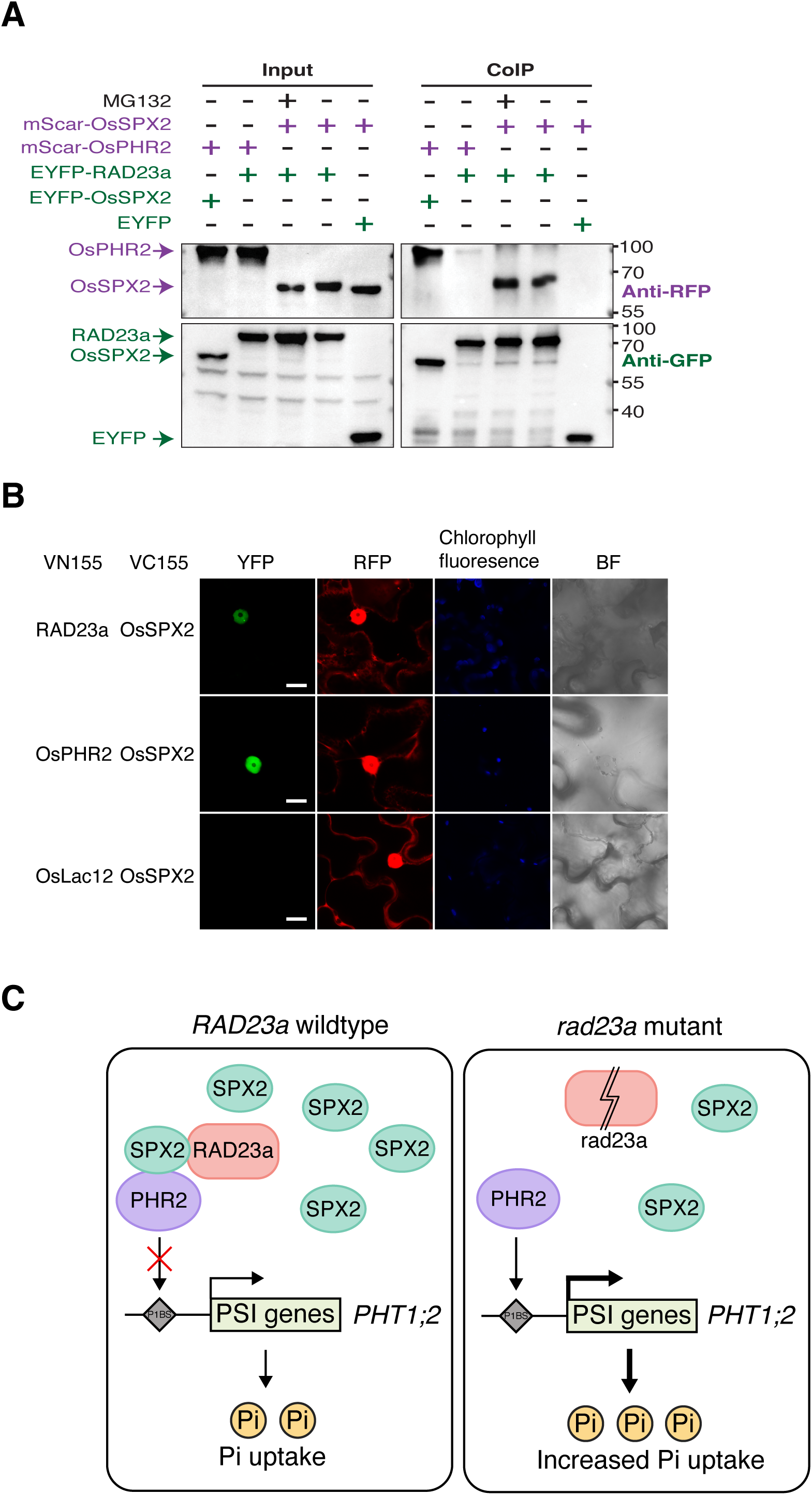
RAD23a interacts with OsSPX2, a negative post-translational regulator of OsPHR2. (A) Co-IP assays showing that RAD23a interacted with OsSPX2 in planta using *N.benthamiana* leaves. RAD23a pulls down mScar-OsSPX2, whereas mScar-OsPHR2 coimmunoprecipitates with EYFP:OsSPX2 but not with EYFP:RAD23a. The +/- symbols indicate protein presence or absence, and protein sizes (kDa) are from the prestained ladder and are provided on the right side of the blot. (B) BiFC assays in epidermal cells of N.benthamiana leaves, showing the interaction of RAD23a and OsSPX2, as well as interaction of OsPHR2 and OsSPX2 (positive control), in the nucleus. Scale bar=20 μm. (C) Proposed working model of RAD23a regulating the Pi uptake. We hypothesize that RAD23a interacts with OsSPX2 directly in a complex involving RAD23a-OsSPX2-OsPHR2. In rad23a mutants, induced edits and/or lower abundance of RAD23a transcript perturbs this complex, resulting in release of additional PHR2 for downstream gene regulation. More available (free) OsPHR2 binds to phosphate transporter (OsPHT1;2) promoters leading to higher Pi uptake in the mutants. *OsSPX2* expression is shown based on the mRNA expression.

To further confirm the in-planta interaction between RAD23a and OsSPX2, we used the BiFC assay by fusing the split halves of mVenus to RAD23a and OsSPX2. As a positive control, PHR2 was co-expressed with OsSPX2 due to their well-established interaction, while rice Laccase with OsSPX2 served as a negative control. Co-expression of RAD23a and OsSPX2 produced nuclear fluorescence, confirming their interaction. Notably, RAD23a and OsSPX2 exhibit a punctate signal, perhaps due to their co-localization in membraneless nuclear condensates (Figure 7B; Supporting Figure S9). We also observed the expected interaction between OsPHR2 and OsSPX2. In contrast, no mVenus fluorescence signal was detected in the negative control, despite the presence of the RFP signal, which served as a transformation and expression control (Figure 7B). These findings suggest that OsSPX2 likely interacts with RAD23a in the nucleus.

## Discussion

In this study, we characterized a rice *Radiation Sensitive23a* (*RAD23a*), a candidate gene identified through GWAS that prioritized causal pre-mRNA splicing events associated with shoot growth and Na^+^ accumulation under saline conditions (Yu *et al*., 2021). In rice, RAD23 family includes four members, which are associated with nucleotide excision repair (NER) and protein degradation through transfer of ubiquitinated proteins to the 26S proteasome (Grønbæk-Thygesen *et al*., 2023; Wang *et al*., 2017). Disruption of *RAD23a* by CRISPR/Cas9 resulted in enhanced shoot growth under control and salt stress (Figure 1B). Furthermore, under saline conditions, Na concentration in shoots and roots were lower in some mutants compared with WT (Figure 2A). For instance, Na concentrations in mut2 and mut3 were significantly lower than that in WT in roots while mut1 were comparable with WT, suggesting that there is tissue-specific variation in Na concentration that is independent of the domain being disrupted by the edits. These results indicate that *RAD23a* negatively regulates the salt-tolerance response in rice at both osmotic (growth) and ionic level (Na uptake). This also supports the previous hypothesis for the role of natural splice variants at the *RAD23a* locus contributing to shoot-level salt tolerance in rice.

The Transcriptome analysis identified that *OsMADS56*, and several dehydration-related genes (*OsOSCA1.1*, *Os04g48230*, *Os12g43720*, *Os07g46670*) were generally upregulated in mutants relative to WT under salt stress (Figure 4B). Cui et al. (2024) recently reported that *OsMADS56* is a causal gene of salt tolerance, with its altered expression resulting from the gene’s presence-absence variations. Our analysis of an RNA-seq dataset for RDP1 genotypes identified that the A-allele genotypes (of *RAD23a*) showed nearly no read counts while the G-allele genotypes exhibited over hundred-fold higher expression on average (Supporting Figure S6). However, the difference in *OsMADS56* expression between WT and mutants was not as pronounced as that observed in allelic variants. It is pertinent to point out that salt stress screening reported by Cui et al. (2024) was conducted under high salt conditions (150 mM NaCl) at germination stages while the *rad23a* mutants’ study was performed under moderate, gradually imposed salt stress (70 mM NaCl) during early vegetative stage. Salt stress sensitivity in rice varies substantially with development stage and the manner of salt stress imposition. Collectively, alteration of salt stress response in *rad23a* mutants could be closely related to Na, K homeostasis, and/or osmotic regulation by multiple dehydration-related genes.

The transcriptome analysis brought forth the unexpected enrichment for phosphorus-related genes in the differentially expressed gene analysis. These transcript-level differences lead to increased inorganic phosphate (Pi) accumulation in shoots of *rad23a* mutants under both control and salt stress, as well as higher root Pi concentration in mutants under salt stress compared to WT (Figure 5B). To investigate the underlying mechanisms, we examined the transcriptome data for Pi-related genes (Figure 5A). Overexpression of *OsPHR2* has been previously shown to upregulate *S-like RNase* genes (*OsRNS4* and *OsRNS5*) in shoots, which play a role in inorganic phosphate (Pi) recycling, leading to increased Pi levels under Pi starvation (Gho *et al*., 2020).

Both *OsRNS4* and *OsRNS5* exhibit shoot-preferred expression in general, with *OsRNS4* exhibiting higher expression levels and *OsRNS5* showing moderate expression. Consistently, the normalized read counts of *OsRNS4* were more than 20-fold higher than those of *OsRNS5* in WT under control conditions. These genes are negatively regulated by *OsPHO2* and strongly induced by salt stress in addition to Pi deficiency (Gho *et al*., 2020; Zheng *et al*., 2014). Our analysis showed that expression of *OsRNS4*, *OsRNS5*, and *OsPHO2* were generally upregulated by salt stress and highly induced in mutants relative to WT (Figure 5A), suggesting the possibility that the repressive effect of *OsPHO2* on *RNS* genes is impaired in *rad23a* mutants through disrupted shuttling of ubiquitinated proteins to the 26S proteasome. However, whether OsPHO2 directly interact with OsRNS4 and OsRNS5 requires further investigation. Notably, expression of *OsTRXh1* generally decreased in mutants relative to WT under salt stress (Figure 5A). *OsTRXh1* is known to be involved in salinity response as well as regulating Pi accumulation (Ying *et al*., 2017; Zhang *et al*., 2011). We also observed allelic difference (A vs G allele) in transcript abundance of *OsRNS4* and *OsTRXh1* was observed under salt stress (Supporting Figure S8). Taken together, the differential regulation of *OsRNS4* and *OsTRXh1* transcript abundance could contribute towards enhanced salt tolerance and increased Pi accumulation in *rad23a* mutants relative to WT.

*OsPHR2* (Rice phosphate starvation response 2) regulates *miR399* and *OsPHT1s* (phosphate transporter 1) expression through binding to the P1BS elements in the promoters of Pi-starvation-induced genes (PSI), including *OsPHT1;2* (Li *et al*., 2015; Liu *et al*., 2010; Zhou *et al*., 2008). *miR399* suppresses the expression of *OsPHO2*, a ubiquitin-conjugating E2 enzyme, which subsequently induces the degradation of several *OsPHT1* family genes under normal conditions (Cao *et al*., 2014; Liu *et al*., 2010; Sun *et al*., 2012; Wu and Wang, 2011). Under both control and salt stress conditions, *OsPHT1;4* was commonly downregulated in mutants relative to WT while *OsPHT1;11* was upregulated specifically in mut1, which exhibits lower transcript abundance of *RAD23a* relative to other two mutants and WT (Figure 5A; Figure 1C). *OsPHT1;11* is involved in symbiotic Pi absorption in rice (Yang et al., 2012) and it rescued a defect in phosphate uptake in a yeast mutant lacking the high affinity phosphate transporter (*pho84*) (Paszkowski *et al*., 2002). Our result suggests that *OsPHT1;11* could be negatively regulated by *RAD23a* transcript abundance in non-symbiotic phosphate responses. It is likely that increased *OsPHT1;11* expression drives the higher Pi uptake in mut1, suggesting that *OsPHT1;11* may have a broader role beyond symbiotic phosphate uptake. OsSPX1, OsSPX2, and OsSPX4 negatively regulate Pi signaling and Pi starvation responses through interacting with OsPHR2 (Lv *et al*., 2014; Wang *et al*., 2014). OsSPX1 and OsSPX2 are exclusively localized in the nucleus whereas OsSPX4 is identified as a membrane-localized protein (Wang *et al*., 2009). Under Pi starvation, the degradation of OsSPX4 in the cytoplasm is accelerated via the 26S proteasome pathway, allowing to release OsPHR2 to the nucleus and activate the expression of PSI genes (Lv *et al*., 2014). Meanwhile, OsSPX1 and OsSPX2 in the nucleus interact with OsPHR2 (Wang *et al*., 2014). Our transcriptome analysis indicates that *OsSPX1*, *OsSPX2*, and *OsSPX4* are downregulated in shoots of *rad23a* mutants, especially under salt stress, whereas WT showed higher expression of these *SPX* genes. Co-immunoprecipitation and BiFC assays in tobacco indicated that RAD23a interacts with OsSPX2, thus providing a mechanistic link between RAD23a and the phosphorus starvation response pathway (Figure 7A&B). Collectively, these results suggest a possible scenario, where OsSPX2 stability is impaired in *rad23a* mutants, or alternatively OsSPX2 and mutated allele’s of RAD23a have a stronger interaction than OsSPX2 and WT allele of RAD23a, subsequently triggering higher OsPHR2 activity. This increased activity due to lower level of available OsSPX2 may enhance the expression of PSI genes, ultimately contributing to the increased Pi levels in mutants.

Similar to the response observed under salinity stress, mutant lines accumulated higher levels of Pi in shoots than WT under Pi-sufficient and low Pi conditions. In shoots, the expression of *OsPHR2* were lower in mutants than WT, and *OsPHO2* were highly induced in mutants under Pi-sufficient conditions. Since *OsPHO2* encodes a ubiquitin-conjugating E2 enzyme that promotes the degradation of *OsPHT1* genes via the 26S proteasome (Cao *et al*., 2014; Liu *et al*., 2010; Sun *et al*., 2012; Wu and Wang, 2011), its upregulation would generally result in reducing Pi uptake. However, despite higher *OsPHO2* expression, the mutants showed increased Pi accumulation and shoot growth. Given that the UBL-UBA protein function includes shuttling ubiquitinated proteins to 26S proteasome (Tsuchiya *et al*., 2017; Zhang *et al*., 2009), it is possible that the edits in *rad23a* mutants may impair the efficient degradation of Pi transporters.

For instance, UBL domain mutant (mut1) may have a reduced shuttling capacity as the RAD23’s UBL domain is known to physically interact with 26S proteasome (Wang *et al*., 2017), resulting in repressed degradation of PHT1 transporters. This is consistent with the observed upregulation of *OsPHT1;2*, *OsPHT1;10*, and *OsPHT1;11* with the strongest induction observed in mut1 than the UBA2 mutants (mut2 and mut3). It is also possible that the increased expression of *OsPHR2* observed in WT is a response to Pi deficiency, which is absent in *rad23a* mutants.

Based on our collective results and previous reports, we hypothesize that RAD23a interacts with OsSPX2 directly in a complex involving RAD23a-OsSPX2-OsPHR2. In *rad23a* mutants, induced edits and/or lower abundance of *RAD23a* transcript perturbs this complex, resulting in release of additional OsPHR2 for downstream gene regulation (Figure 7C). More available (free) OsPHR2 binds to phosphate transporter promoters leading to higher Pi levels in the mutants. Since OsSPX2 transcript abundance decreases in the *rad23a* mutants, it opened the possibility that RAD23a is also involved in transcriptional regulation of OsSPX2 as a second layer of regulating Pi homeostasis by directing a yet unknown transcriptional repressor of OsSPX2 to the 26S proteosome.

Previous reports have linked Pi homeostasis with salt stress response mechanisms in plants (Ahmed *et al*., 2018; Hasanuzzaman and Fujita, 2022; Su *et al*., 2022). For instance, higher Pi accumulation in *Siz1* and *Pho2* mutants in Arabidopsis exhibit increased salt tolerance (Miura *et al*., 2011). *OsPHL7* is another example where phosphate homeostasis and salt stress in rice overlap (Yang *et al*., 2024). Notably, our results contrast from previous findings in that under Pi-sufficient conditions, in *rad23a* mutants, expression of *OsPHO2* is upregulated while *OsPHR2* is downregulated, yet both Pi uptake was increased and shoot growth is enhanced in mutants relative to WT. Collectively, our hypothesis suggests that *RAD23a* may influence Pi transport by modulating protein turnover of Pi transport proteins via shuttling. Therefore, *RAD23a* could regulate salt stress adaptation indirectly through altered Pi accumulation in rice, as well as direct regulation thorough Na, K uptake and osmotic responses associated with dehydration-related genes.

## Author Contributions

H.W. conceived the idea. H.W. and C.Z. designed the experiments. S.O. performed most of the experiments. B.A. preformed protein interaction analysis. J.S.D. generated mutants. S.O. and A.K.N.C analyzed the transcriptome data. S.O. and H.W. wrote the manuscript.

## Acknowledgements

We thank Javier Seravalli from Redox Biology Center at the University of Nebraska-Lincoln for performing for elemental analysis by ICP-MS. We also thank Bara Altartouri for the technical assistance with confocal imaging performed at the UNL Microscopy Core Research Facility (RRID: SCR_017798), supported by the Nebraska Research Initiative and NIH grant P20GM113126.

## Accession numbers

The RNA-seq data are available in the NCBI SRA under BioProject ID PRJNA1510219.

## Supporting Figures and Tables

Supporting Figure S1 | Comparison of *RAD23a* gene and protein sequence between the G and the A allele genotypes. The A allele genotypes retain the sixth intron, resulting in a premature stop codon whereas G allele genotypes splice out the sixth intron (Yu *et al*., 2021).

Supporting Figure S2 | Pearson correlation between projected shoot area (PSA) and shoot DW (g) at 27 DAT with confidence interval set at 95% (shaded area).

Supporting Figure S3 | Root DW (g) at 27 DAT under control and salt stress conditions. Statistical significance was determined using two-way ANOVA followed by Duncan’s multiple range test (n=4). Different letters denote significant differences at *P* < 0.05.

Supporting Figure S4 | (A) Potassium concentrations and (B) Na/K ratio in roots and shoots at 27 DAT. Statistical significance was determined using two-way ANOVA followed by Duncan’s multiple range test (n=3-4). Different letters denote significant differences at *P* < 0.05.

Supporting Figure S5 | Relative transcript abundance of several HKT genes at 17 DAT, 7 days after the initial salt stress in roots and shoots. The expression of *OsHKT2;3* wasn’t detected in root tissue. Data represent means from two biological replicates, each with two technical replicates. Error bars represent ± SE. Statistical significance was determined using two-way ANOVA followed by Duncan’s multiple range test. Different letters denote significant differences at *P* < 0.05.

Supporting Figure S6 | Allelic difference in shoot transcript abundance (normalized read counts) of several salt- and/or dehydration-related DEGs identified in this study. The A allele (n=11) and the G allele (n=90) genotypes were compared using two-way ANOVA followed by Duncan’s multiple range test. Error bars represent ± SE and different letters denote significant differences at *P* < 0.05. The raw transcriptomic data can be found in NCBI GEO database (Accession #: GSE98455) from which *RAD23a* was identified (Du *et al*., 2019; Yu *et al*., 2021).

Supporting Figure S7 | Allelic variation in *RAD23a* transcript abundance in 7-day old seedling leaf tissues among three G allele accessions and two A allele accessions. The corresponding seeds were germinated and grown in half-strength MS agar. Data represent means from two biological replicates, each with three technical replicates. Error bars represent ± SD. The 5 accessions are NSFTV-70, NSFTV-379, Kitaake, NSTFV-117 and NSFTV-78 from rice diversity panel 1. The dotted lines represent average of transcript abundance of *RAD23a* within allelic accessions. Student’s t-test was used to compare the allelic variations (****, *P*-value < 0.0001).

Supporting Figure S8 | Allelic difference in shoot transcript abundance (normalized read counts) of several Pi-related DEGs identified in this study. The A allele (n=11) and the G allele (n=90) genotypes were compared using two-way ANOVA followed by Duncan’s multiple range test. Error bars represent ± SE and different letters denote significant differences at *P* < 0.05. The raw transcriptomic data can be found in NCBI GEO database (Accession #: GSE98455) from which *RAD23a* was identified (Du *et al*., 2019; Yu *et al*., 2021).

Supporting Figure S9 | BiFC assays in epidermal cells of *N.benthamiana* leaves, showing the interaction of RAD23a and OsSPX2 in the nucleus. Multiple independent infiltration sites in different leaves were imaged as biological replicates. Scale bar=20 μm.

Supporting Table S1 | List of primers used in this study.

Supporting Table S2 | List of candidates tested for yeast two-hybrid (Y2H) assays.

Supporting Table S3 | Coding sequences and codon-optimized sequences used in the interaction studies.

Supporting Table S4 | RNA-seq normalized read counts of salt- and/or dehydration-related genes in WT and *rad23a* mutants under control and salt stress.

