## Supporting figures for "*OsRAD23a* negatively regulates salt tolerance and phosphorus uptake in rice"

### Supplementary Figure S1

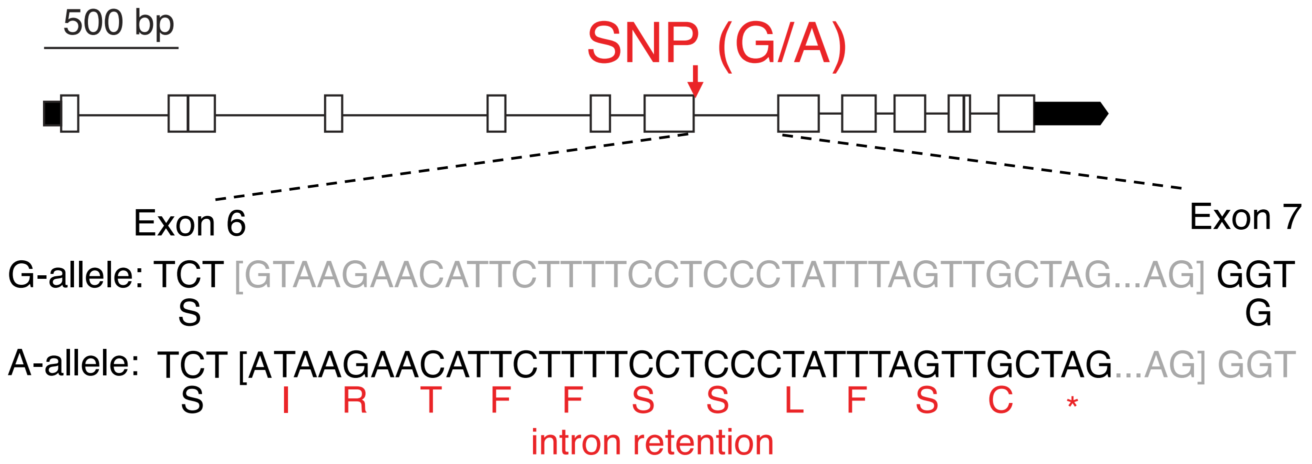


### Supplementary Figure S2

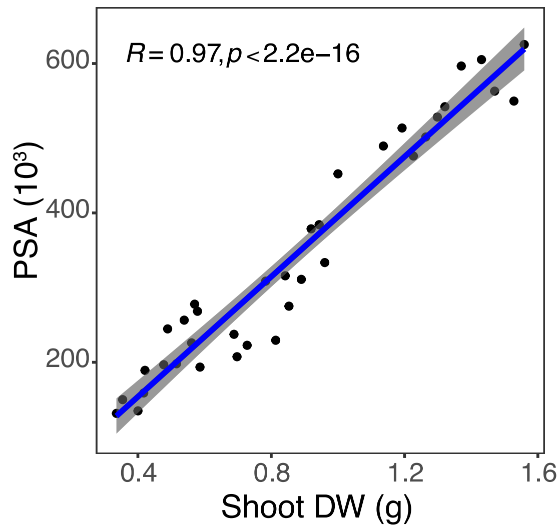


### Supplementary Figure S3

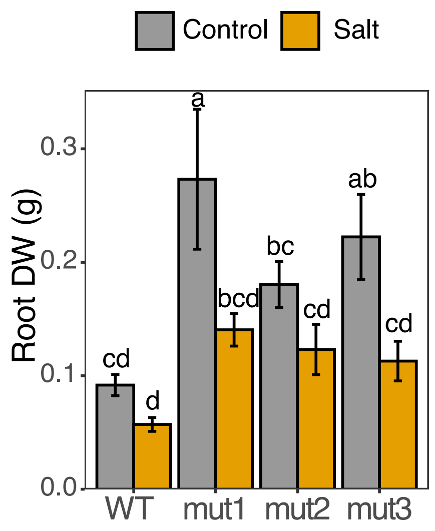

### Supplementary Figure S4

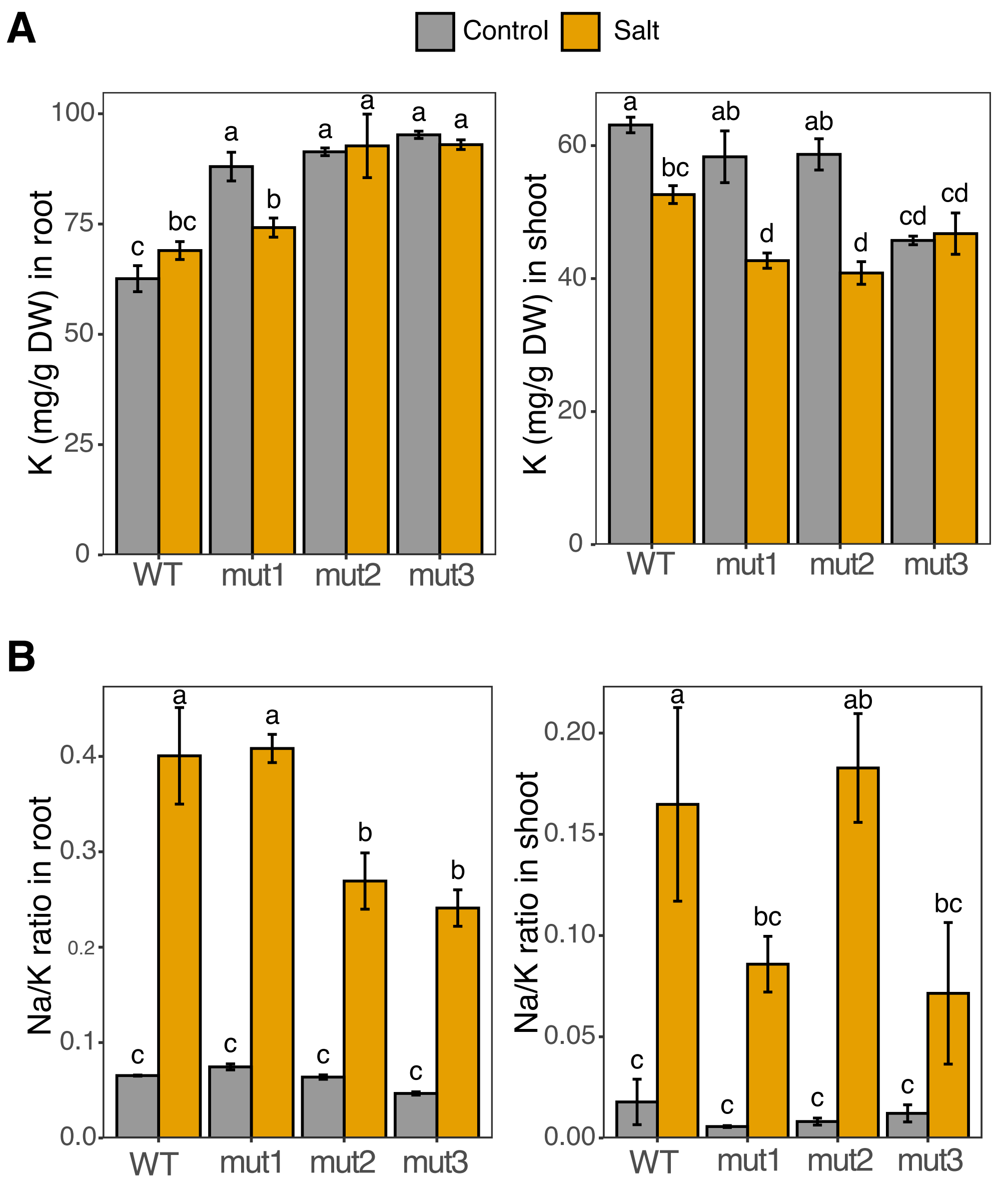

### Supplementary Figure S5

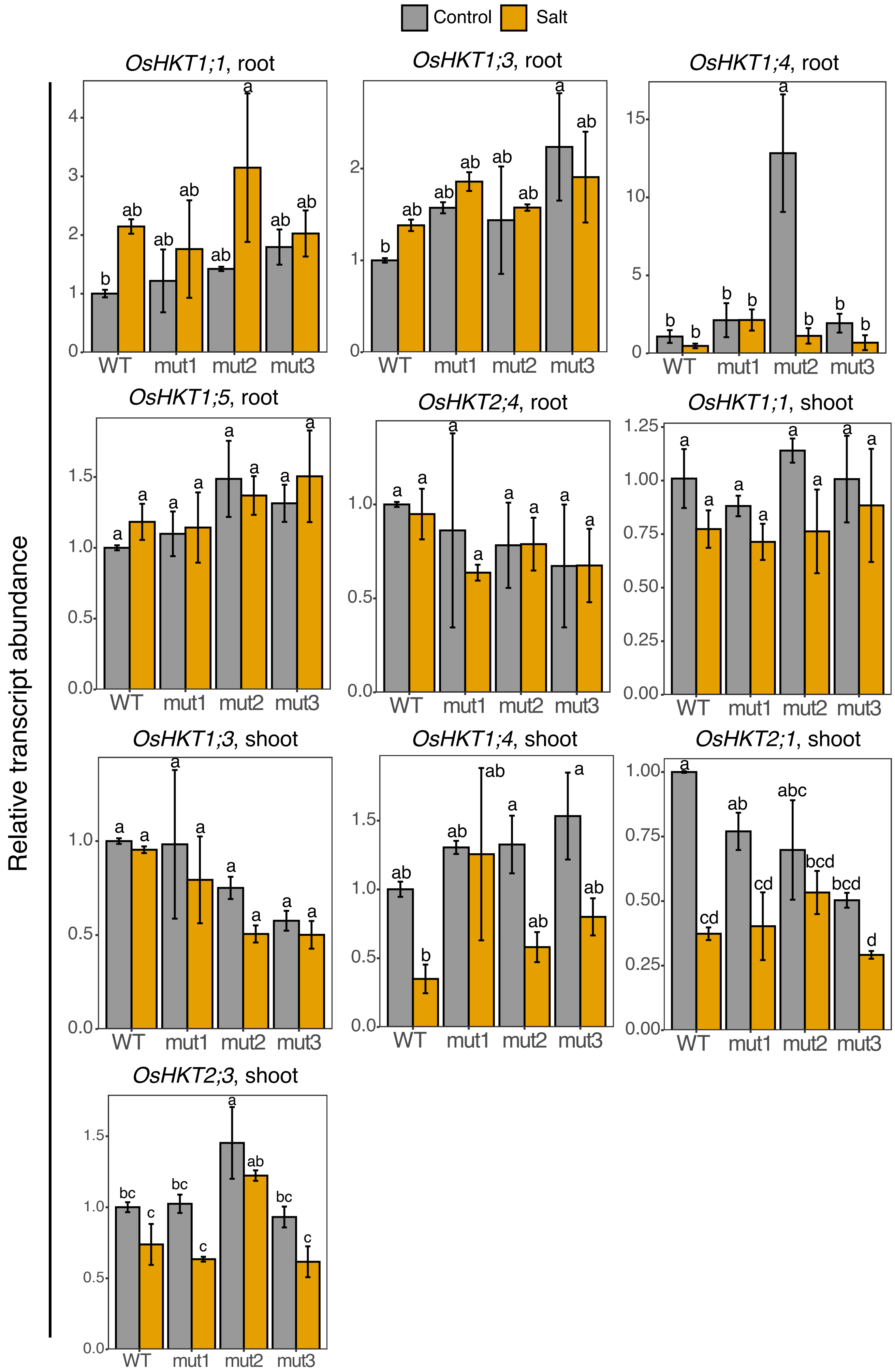

### Supplementary Figure S6

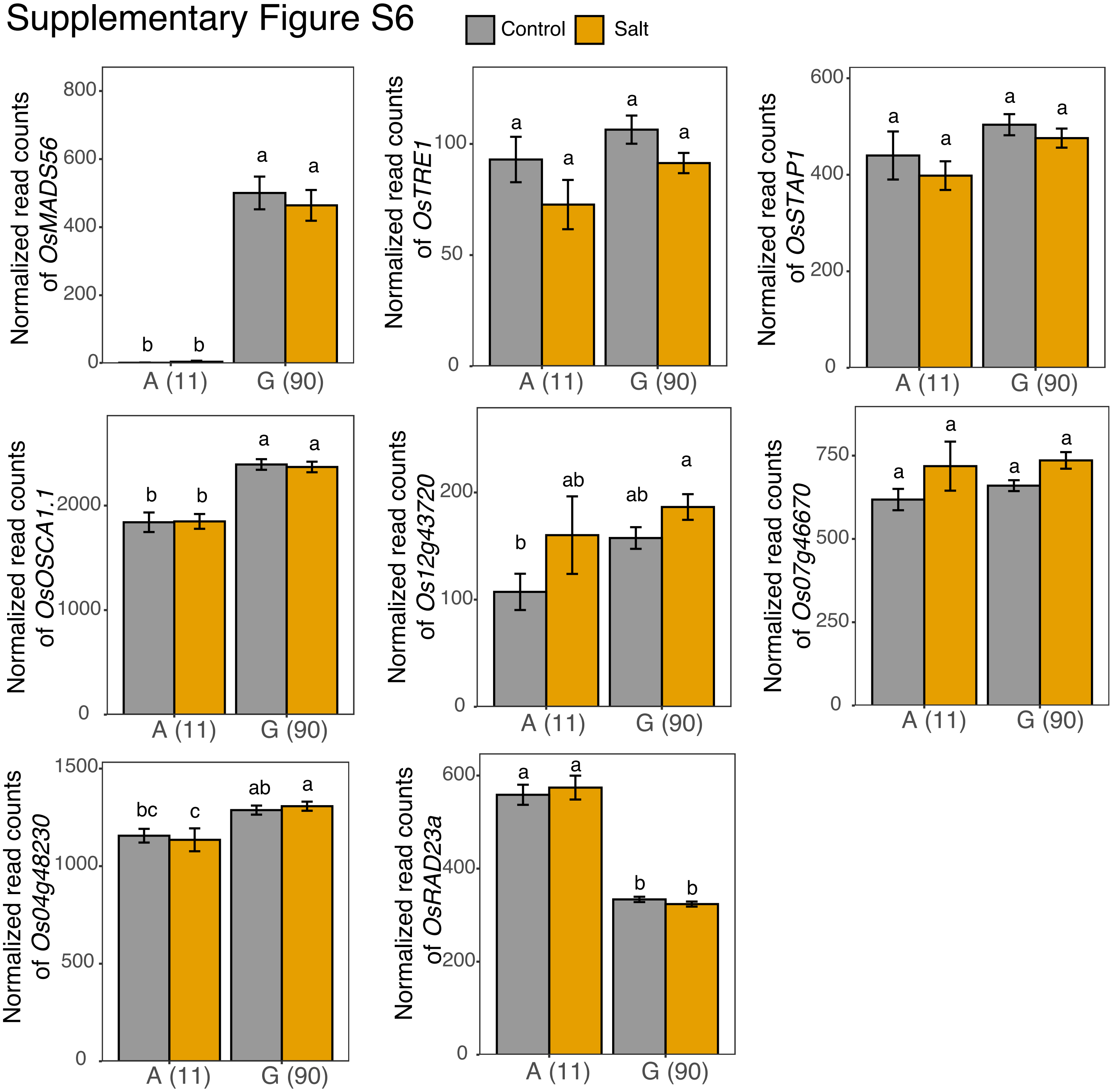


### Supplementary Figure S7

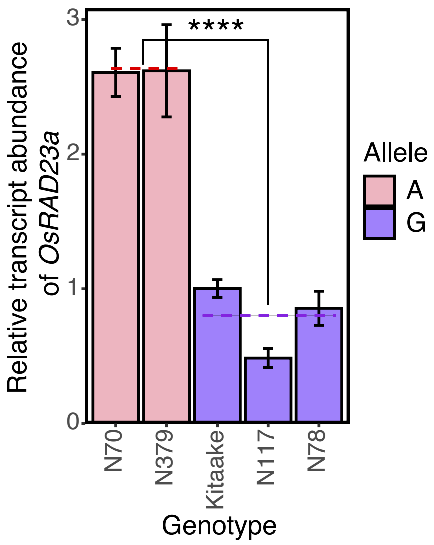

Supplementary Figure S8

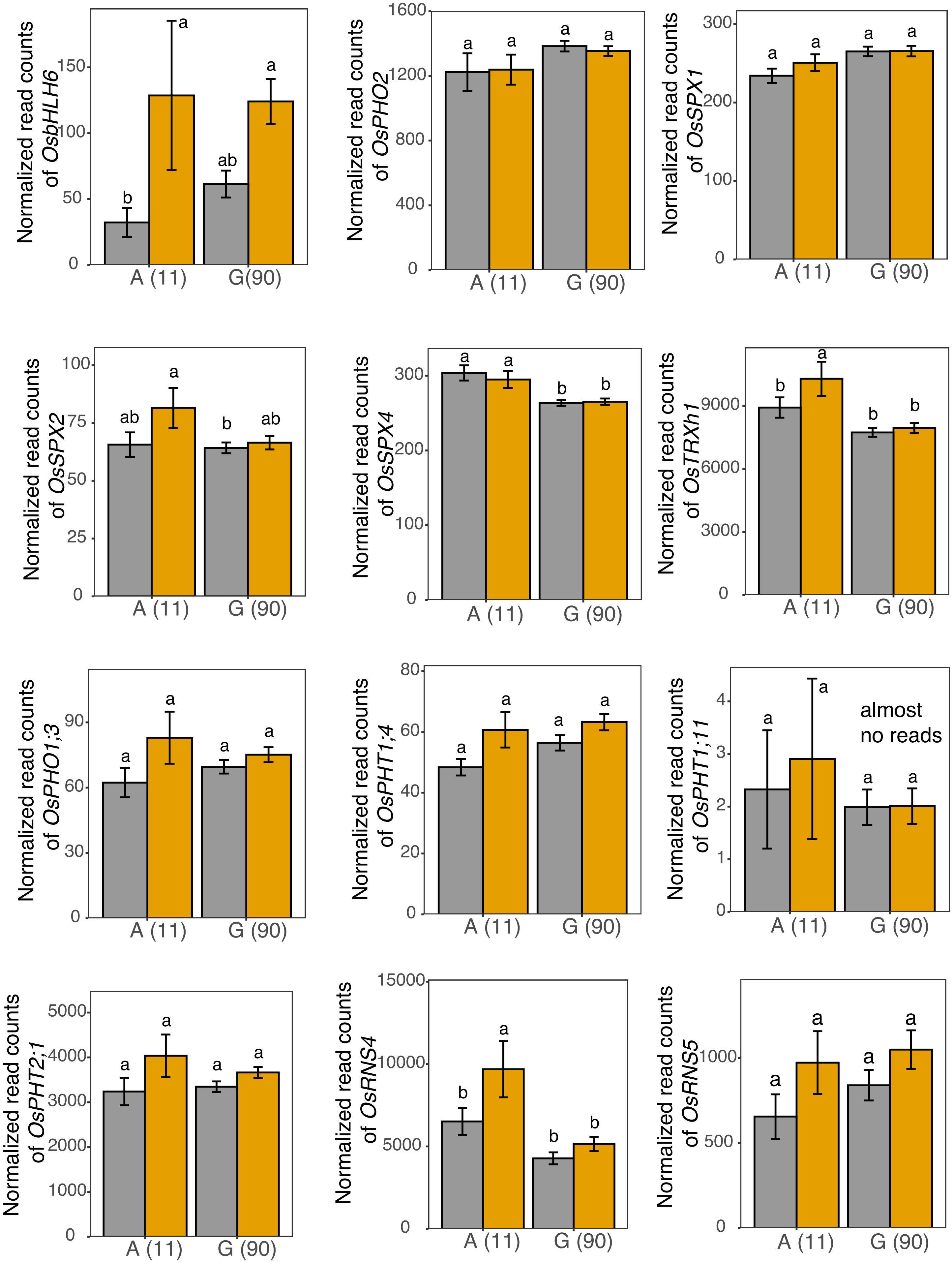

### Supplementary Figure S9

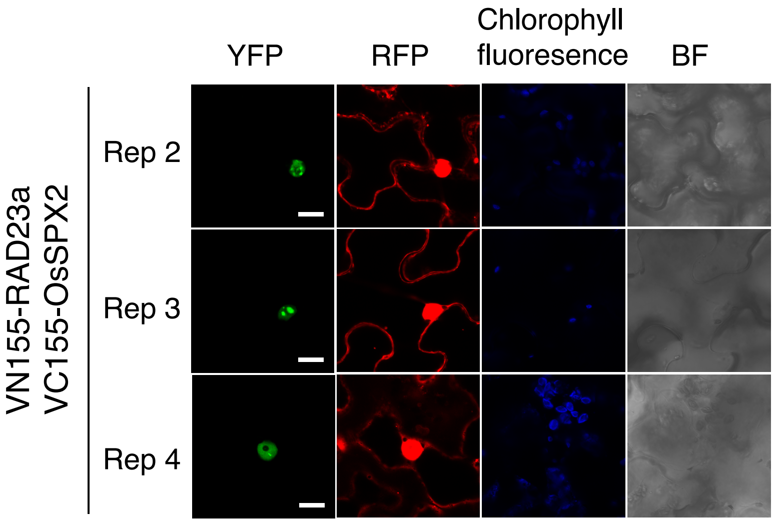
